# FOXP3-engineered regulatory T cells restore intestinal barrier integrity in Crohn’s disease enteroids via PDGF-AA

**DOI:** 10.64898/2026.05.25.727767

**Authors:** Neetu Saini, Babajide A. Ojo, Niharika Bozza, Akshaya Ramachandran, Pinki Gahlot, Javier A. López-Rivera, Tracy Tran, Peacha Sokzini, Hemmo Meyer, James C.Y. Dunn, Maria Grazia Roncarolo, Michael J Rosen, Rosa Bacchetta

## Abstract

Epithelial regeneration and barrier integrity are impaired in inflammatory bowel diseases, including Crohn’s disease (CD), yet current therapies largely target immune inflammation without directly promoting mucosal repair. While regulatory T cells are classically immunomodulatory, their capacity to directly support human intestinal stem cells (ISCs) and barrier function remains unclear. In this study, we tested the hypothesis that FOXP3-expressing regulatory T cells—engineered CD4^LVFOXP3^ and thymic-derived T_reg_ (tT_reg_)—directly support human ISC maintenance and restore epithelial barrier function independent of their immunomodulatory function. Using CD patient ISCs-derived enteroids that display disease-associated damage, we established co-culture with FOXP3-engineered T_reg_ cell-CD4^LVFOXP3^ or thymic-derived T_reg_ (tT_reg_). The presence of either CD4^LVFOXP3^ or tT_reg_ cells enhanced enteroid growth, improved epithelial barrier function, and restored apical-basal polarity of ISCs, indicating reparative capacity. Conversely, activated conventional CD4^+^ T cells reduced barrier function and abrogated apical-basal polarity. Integrating secretome profiling with ligand add-back and receptor or ligand blockade, we identify the PDGF-AA–PDGFRα axis as a key regulator of T_reg_-mediated intestinal epithelial barrier integrity, but dispensable for T_reg_ suppressive capacity. Collectively, our data delineate a direct, human tissue-intrinsic role of FOXP3-driven Treg in the interaction with ISCs via PDGF-AA-PDGFRα, enhancing epithelial barrier function and positioning CD4^LVFOXP3^ as a treatment approach coupling immunoregulation with epithelial repair.

**One Sentence Summary:** Regulatory T cells improve the intestinal epithelial barrier function, primarily, through the PDGF-AA–PDGFRα axis

## INTRODUCTION

Regulatory T (T_reg_) cells have long been recognized as the primary mediators of immune regulation. Their therapeutic potential is fundamentally rooted in their ability to suppress effector T-cell function, downregulate antigen presentation, and dampen inflammatory cascades to maintain a tolerogenic environment (*1, 2*). These functions are governed by the constitutive high expression of the transcription factor FOXP3, which drives the gene programs required for lineage stability and suppressive capacity (*3, 4*). Crucially, the functional efficacy of T_reg_ cells depends upon TCR-mediated activation within tissue-specific contexts, allowing for targeted modulation of the immune landscape (*5, 6*). In recent years, the function of T_reg_ cells has expanded beyond canonical immune suppression and they have been implicated as regulators of tissue integrity and regeneration via specific mechanisms (*7, 8*). This pro-regenerative capacity, which is exerted in a TCR-independent fashion, has been documented, in the skin, lung, thymus and skeletal muscle. Thus, it has been proposed that T_reg_ cells facilitate repair through two distinct pathways: an indirect mechanism which limits immunopathology and bystander damage through the control of T cell mediated damage; and a direct mechanism, involving the secretion of molecules to promote repair

While murine studies have identified several mediators of T_reg_-driven gut repair—including Fibroblast Growth Factor 2 (FGF2), IL-17, and IL-6R trans-signaling (*9, 10*)—mechanistic insights into human T_reg_ contributions are sparse. Current understanding of human T cell–mediated epithelial repair is primarily limited to Type 1 regulatory T (Tr1) cells, which via IL-10 decrease inflammation, and promote goblet cell differentiation and barrier function, possibly via, IL-22 signaling (*11*). Whether human thymic-derived T_reg_ (tT_reg)_ cells interact with the intestinal epithelium and modulate intestinal stem cell (ISC) niches remains a major gap in our understanding of immune–epithelial crosstalk. A primary reason for the knowledge gap regarding human T_reg_ cell function in the intestinal tissue, is the technical challenge of isolating T_reg_ in sufficient numbers from patient blood or/and a lack of suitable humanized mouse model to explore human Tregs-intestinal epithelial interaction and repair to conduct robust functional studies.

Our laboratory has engineered T_reg_ cells (CD4^LVFOXP3^), which can be generated in large quantities while maintaining their suppressive and tT_reg_ phenotypic identity. CD4^LVFOXP3^, generated by lentiviral delivery of FOXP3 into total CD4^+^ T cells (*12, 13*), phenotypically and functionally resemble tT_reg_ cells and are currently in Phase I clinical trial (NCT05241444), as a T_reg_ cells replacement therapy for Immune dysregulation, Polyendocrinopathy, Enteropathy, and X-linked (IPEX) syndrome, all caused by FOXP3 mutations. In this pediatric monogenic autoimmune disease, the primary symptom is severe enteropathy, resembling severe Crohn’s disease (CD), and IPEX is classified as a very-early onset (VEO)-IBD. This Tregopathy emphasizes the critical role of regulatory T cells (Tregs) in maintaining gut homeostasis (*14*). In addition, multiple studies have demonstrated both qualitative and quantitative defects in Tregs in the pathogenesis of CD (*14, 15*). These findings provide a strong rationale for the development of Treg-based therapies, several of which are currently being evaluated in early-phase clinical trials for inflammatory bowel disease (IBD) (*1, 15*).

We employed ISCs derived human enteroids from CD patients—an *in vitro* system that preserves disease-specific epithelial traits and recapitulates the complex architecture, barrier function, and regenerative dynamics of the intestinal epithelium. Utilizing autologous/allogenic co-culture of activated CD4+ T cells and enteroids derived from patients with CD, we established a model of damaged intestinal epithelium to investigate the function of T_reg_ cells. Using this system, we demonstrate that T_reg_ cells interact directly with ISCs to drive their regenerative capacity resulting in significantly enhanced barrier function via increased enteroid growth and improved apical-basal polarity of the cells. Using neutralizing antibody and pharmacological inhibition, we identify the Platelet-derived growth factor-AA (PDGF-AA) signaling pathway as a central mechanism underlying this interaction. While the PDGF family members are known regulators of mesenchymal and stromal cell activity, they have not previously been identified as critical mediators of tT_reg_-ISC interaction. Our data reveal that T_reg_-derived PDGF-AA acts directly on the ISCs via - Platelet-derived growth factor receptor alpha (PDGFRα), offering a novel therapeutic mechanism, supporting the use of T_reg_ cell therapies for restoring intestinal homeostasis in IBD, not only by dampening inflammation and suppressing pathogenic effector T-cells but also promoting tissue repair and healing. This would make T_reg_ cell therapies superior to any current pharmacological treatment.

## RESULTS

### Co-culture of autologous CD4^LVFOXP3^ promotes the growth of CD patient-derived enteroids

We have extensively characterized the CD4^LVFOXP3^ cells generated from healthy donors (HD) and shown that they have overlapping tT_reg_ phenotypes and functions (*12, 13, 16*). CD4^+^ T-cell subsets in peripheral blood from CD patients show altered naïve/memory distributions and increased circulating TH17 cells (*17–19*). To determine whether these disease-associated differences in the starting CD4^+^ T-cell pool influence the generation of engineered CD4^LVFOXP3^ T_reg_ cells, we first compared transduction efficiency, FOXP3 expression, T_reg_-associated marker expression, and suppressive function of CD4^LVFOXP3^ cells generated from CD patients with active disease or in remission (Table S1, **fig. S1A**). Transduction efficiency, assessed by the surface expression of the truncated Nerve Growth Factor (NGFR) included in the LVFOXP3 construct, was comparable between cells derived from individuals with CD and HD (**fig. S1B**). Regardless of disease status, as expected, the CD4^LVFOXP3^ expressed FOXP3 and NGFR (**fig. S1C, D**), and showed high frequency of the tT_reg_ cells surface markers (CD127^low^, CD25^+^), and no difference was observed between CD–derived and HD–derived CD4^LVFOXP3^ cells (**fig. S1E**). Expression of additional tT_reg_ cell-associated markers, including Helios, GITR, PD-1, ICOS, and CTLA-4, and cytokine receptors (e.g., IL-1R and IL-6R) was also similar between CD- and HD-derived CD4^LVFOXP3^ cells, (**fig. S1F**). CD4^LVFOXP3^ cells derived from individuals with CD suppressed the proliferation of responder allogenic CD4^+^ T cells, whereas NGFR-only control cells (CD4^NGFR^) did not (**fig. S1G, H**). The magnitude of suppression by CD–derived CD4^LVFOXP3^ cells was comparable to that observed in HD-derived CD4^LVFOXP3^ cells (**fig. S1H**). Collectively, these data indicate that CD–derived CD4^LVFOXP3^ cells recapitulate the expression of key markers associated with tT_reg_ and suppressive functions to an extent comparable to HD-derived CD4^LVFOXP3^ cells, supporting their use in this study as a surrogate tT_reg_ cell for investigating the interactions with ISCs.

To this aim, we first generated and cultured ISCs-enriched enteroids starting from intestinal crypts isolated from ileal biopsies of CD patients or HD. Crypts were embedded in Matrigel and cultured in organoid growth complete medium (OGM-CM) containing key niche factors and grown into three-dimensional epithelial spheroids (enteroids) as shown (**Fig. 1A)**. Undifferentiated enteroids obtained from CD patients (n=5) were enriched for LGR5^+^ ISCs and LGR5 expression levels were comparable between CD- and HD–derived enteroids (**Fig. 1B, C**). Enteroid size, measured as area, was also similar between enteroids derived from CD- and HD-derived cells (**Fig. 1D**).

**Figure 1:**
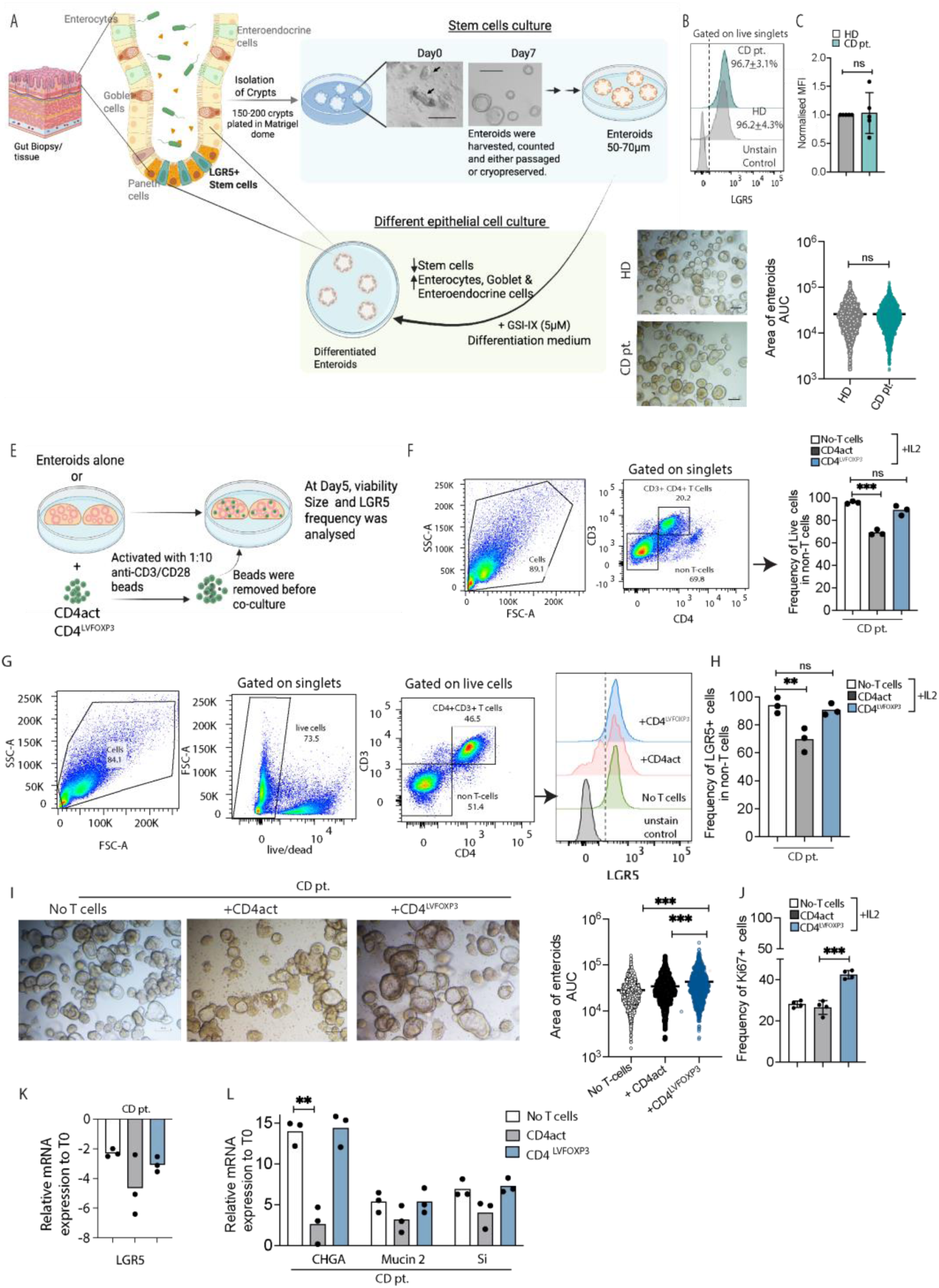
CD4^LVFOXP3^ maintains LGR5^+^ cells and promotes the growth of CD patient-derived enteroids. **(A)** Schematic of intestinal crypt isolation and enteroid culture from ileal tissue or biopsies. **(B, C)** LGR5 expression in enteroid cells assessed by flow cytometry: frequency of LGR5⁺ cells in enteroid from CD patients and HD (B), and fold change in LGR5 MFI in enteroid cells from CD patients, relative to HD (C). **(D)** Representative brightfield images and size quantification of CD patients and HD-derived enteroids. **(E)** Schematic showing co-culture of enteroids with activated CD4^LVFOXP3^ or CD4act (1:100). (**F)** Frequency of live cells in non-T cells in enteroids co-cultured with activated CD4^LVFOXP3^ or CD4act. **(G, H)** Flow cytometric analysis of LGR5 expression in live non-T cells, including representative histograms (G) and quantification (H). **(I, J)** Brightfield images and size quantification of CD-derived enteroids **(I),** and frequency of the Ki67^+^ cells within LGR5^+^ cells in enteroids **(J)**, cultured alone or with indicated T cells. (**K, L)** Relative expression of LGR5 (J) and differentiation markers Chromogranin A (CHGA), Sucrose-isomaltase (SI), and Mucin 2 (K) in enteroids cultured alone or with indicated autologous T cells in ODM-CM. Data are from five (B) and three independent (C-K) CD patients. Welch’s t-test (B–C) and paired t-test (I–K); **P < 0.01, ***P < 0.001; ns, not significant.

To assess the impact of CD4^LVFOXP3^ on the undifferentiated enteroids, we co-cultured them with autologous CD4^LVFOXP3^ cells or control untransduced CD4^+^ T cells (CD4act) pre-activated with anti-CD3/CD28 dynabeads (**Fig. 1E**). Following the co-culture (**Fig. 1E**), the frequency of the live cells was maintained ∼89% in enteroids co-cultured with CD4^LVFOXP3^ as in the enteroids cultured without T cells. In contrast, enteroid showed a significantly reduced proportion (∼69%) of live cells in co-culture with autologous CD4act (**Fig. 1F**). These results indicate that CD4^LVFOXP3^ cells do not induce cell death of LGR5^+^ ISCs, whereas CD4act-reduced numbers of viable ISCs. In addition, the frequency of LGR5^+^ cells (∼90%) was maintained in enteroids co-cultured with CD4^LVFOXP3^ (**Fig. 1G, H)** but not with autologous CD4act T cells (∼69%). (**Fig. 1G, H**). Moreover, the average size of the enteroids co-cultured with CD4^LVFOXP3^ significantly increased (p-values <0.001) compared to enteroids cultured alone or with CD4act (**Fig. 1I**). The size of the LGR5^+^ cells weas comparable between enteroids cultured alone or co-cultured with CD4^LVFOXP3^ or CD4act (**fig. S2A**). The frequency of proliferating LGR5^+^ cells was significantly (p<0.001) increased in enteroids co-cultured with the CD4^LVFOXP3^ as compared to those cultured without T cells or CD4act (**Fig. 1J, fig. S2B**). Together, these data suggest that CD4^LVFOXP3^ cells promote enteroid growth by enhancing LGR5^+^ cell proliferation.

To determine whether CD4^LVFOXP3^ affects LGR5^+^ ISC differentiation, we induced differentiation by culturing enteroids in organoid differentiation complete medium (ODM-CM) and quantified the expression of several distinct epithelial lineage marker transcripts, Chromogranin A (enteroendocrine cells), Mucin 2 (MUC2; goblet cells), Sucrase Isomaltase (SI; enterocyte cells), and LGR5 (ISCs). As expected, ODM-CM reduced LGR5 expression regardless of co-culture condition (**Fig. 1K**), consistent with the initiation of epithelial stem cell differentiation. In the presence of CD4^LVOFXP3^, we observed comparable induction of Chromogranin A, MUC2, and SI (**Fig. 1L**), indicating their preservation of differentiation capacity. In contrast, co-culture with autologous CD4act diminished the induction of these lineage markers, significantly for Chromogranin A (p value=0.002), indicating impaired differentiation (**Fig. 1L**). Taken together, the co-culture data indicate that CD4^LVFOXP3^ cells do not adversely affect LGR5^+^ stem-cell integrity and differentiation, whereas CD4act display pathogenic effector T-cell activity, causing loss of LGR5^+^ cells, and impaired differentiation that was more evident for enteroendocrine cells.

### CD4^LVFOXP3^ or tT_reg_ cells enhance the epithelial TEER and reduce polarity defects

One of the primary functions of intestinal epithelium is to act as a barrier between the luminal and the basolateral side to prevent the movement of pathogens and antigens (*20*). Barrier defects are a hallmark of IBD, including CD (*21*). We therefore determined whether this barrier alteration is recapitulated in enteroid-derived monolayers from CD patient’s vs HD. Transepithelial electrical resistance (TEER) across the enteroid monolayer was measured to examine barrier function. A higher TEER indicates strong intracellular tight junctions between cells, resulting in the high resistance corresponding to enhanced barrier function, while lower resistance denotes leaky epithelium. Results showed that the monolayers derived from CD patient enteroids had significantly lower (p-value= 0.017) TEER, compared with monolayers derived from HD, which was evidence of a leaky epithelial barrier in CD (**Fig. 2A, B**). Co-culture with autologous CD4^LVFOXP3^, seeded in the bottom well, significantly increased TEER compared to CD monolayers cultured alone (**Fig. 2C**). In contrast, co-culture with autologous CD4act cells reduced TEER relative to monolayers cultured without T cells, indicating further disruption of epithelial barrier function (**Fig. 2C**). Extending these findings, co-culture of CD patient-derived epithelial monolayers with allogeneic tT_reg_ cells, CD4^LVFOXP3^ cells, or autologous CD4^LVFOXP3^ cells yielded comparable, significant increases in TEER **(Fig. 2D**), indicating that CD4^LVFOXP3^ cells replicate the barrier-protective activity of tT_reg_ cells. To assess effects on mature epithelium, confluent epithelial monolayers were switched to differentiation medium, and autologous or allogeneic CD4^LVFOXP3^ cells or allogeneic tT_reg_ cells were activated basolaterally (**Fig. 2E**). Differentiated monolayers from CD enteroids exhibited reduced TEER relative to HD, recapitulating the barrier defect, and co-culture with CD4^LVFOXP3^ cells (autologous or allogeneic) or tT_reg_ cells significantly increased TEER **(Fig. 2F).** Importantly, analysis of differentiation in these enteroid monolayers revealed comparable composition of distinct epithelial cell types between HD- and CD-derived cultures. Furthermore, co-culture with either autologous or allogeneic CD4^LVFOXP3^ or tT_reg_ cells did not alter epithelial cell type composition (**fig. S3A, B**).

**Figure 2:**
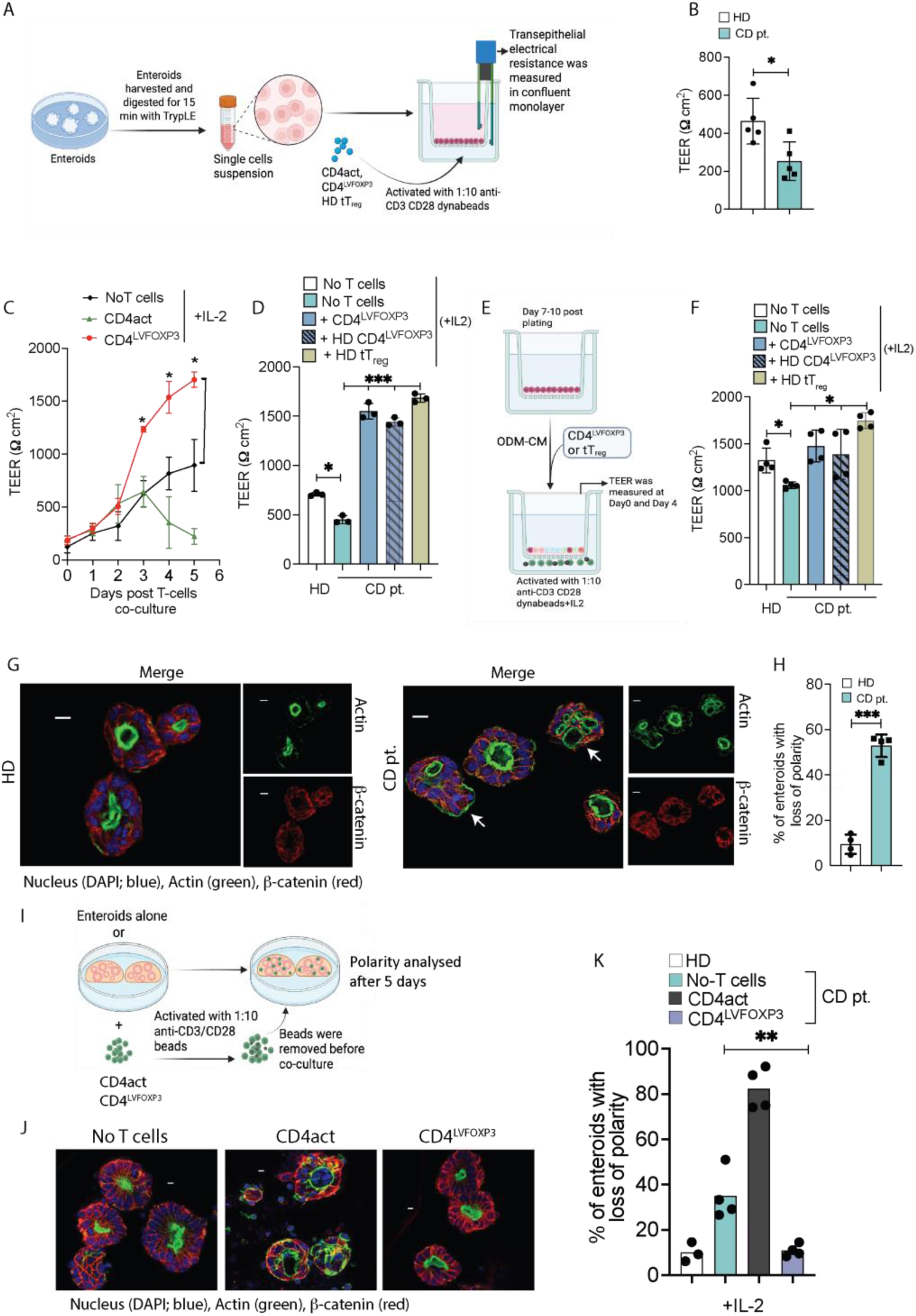
CD4^LVFOXP3^ improves TEER and cell polarity. **(A)** Schematic showing the setup for TEER analysis and co-culture with CD4^LVFOXP3^ or tT_reg_ cells. **(B-D)** TEER measurements in monolayer derived from CD patients or HD enteroids (B), or monolayers co-cultured with the autologous CD4act or CD4^LVFOXP3^ cells (C), or with allogenic CD4^LVFOXP3^ or tT_reg_ cells from HD (D). **(E,F)** Schematic showing the setup for TEER analysis in differentiated monolayer co-culture with CD4^LVFOXP3^ or tT_reg_ cells (E) and TEER values in monolayers co-cultured in these conditions (F). **(G)** Representative confocal images of HD- and CD-derived enteroids stained for F-actin (phalloidin–Alexa 488), β-catenin, and nuclei (DAPI); arrowheads indicate misoriented cells. **(H)** Percentage of enteroids with misoriented cells. **(I–K)** Schematic of enteroid co-culture with preactivated CD4act or CD4^LVFOXP3^ T cells (I); representative images stained as in G (J); quantification of enteroids with misoriented cells in the indicated conditions (K). Data plotted are independent experiments performed with five (B) or four (F-J) or three (C, D) CD patients or HD. Statistical analyses were performed using Welch’s t-test (B, H) and paired t-test (C-F, J). ** and *** denote p < 0.01 and p < 0.001, respectively; ns denotes p > 0.05.

Tight junctions govern epithelial barrier function and establish apical–basolateral polarity of the epithelial cells (*22–24*); this polarity is preserved during enteroid culture. In HD-derived enteroids, F-actin lined the luminal (apical) surface and β-catenin marked the outer basolateral membranes, consistent with preserved polarity (**Fig. 2G**). In contrast, we observed that a subset of CD-derived enteroids contained cells with aberrant orientation (arrowheads) and were scored as misoriented enteroids (**Fig. 2G, fig. S3C)**. Analysis of polarity revealed that enteroids derived from CD exhibited a significantly increased numbers of misoriented enteroids, compared to those derived from HD (**Fig. 2H**). Notably, the addition of CD4^LVFOXP3^ significantly reduced the number of misoriented enteroids, whereas CD4act increased this fraction (**Fig. 2I-K**).

Taken together, these data indicate that CD4^LVFOXP3^ cells enhance barrier function and rescue polarity defects in CD-derived enteroids, whereas CD4act cells further increased barrier and polarity defects. The physical separation of epithelium and T cells in these experiments suggests the potential existence of paracrine mediators, which we examined in subsequent experiments.

### CD4^LVFOXP3^ or tT_reg_ cells reduce the secretion of pro-inflammatory cytokines and increase growth factor levels in co-culture supernatants

To assess the potential paracrine mediators of T-cell-epithelial crosstalk, supernatants were harvested from enteroid monolayers co-cultured with or without T cells from CD patients (n=5) and 80 soluble mediators, including pro- and anti-inflammatory cytokines, chemokines, growth factors, colony-stimulating factors, and apoptotic ligands, were quantified by Luminex. Only a few epithelial-derived cytokines, including IL-7, CCL2 and EOTAXIN2, were detected in supernatants from enteroid monolayers cultured without T cells (**Fig. 3A**). Many cytokines and growth factors were detected in the supernatants from the co-culture of enteroids with CD4^LVFOXP3^ cells or CD4act (**Fig. 3A, fig. S4A, B).** Relative to those from CD4act co-cultures, supernatants from CD4^LVFOXP3^ co-cultures contained significantly lower concentrations of Fas ligand (FasL), Interferon-gamma (IFN-γ), Th2-associated cytokines (IL-13, IL-5, IL-9 and IL-3), the chemokine CCL1, and soluble mediators linked to epithelial injury (sCD40L, sICAM-1 and IL-1α) (**Fig. 3B, fig. S4B**). Pro-inflammatory cytokines, including IFN-γ, IL-13, were previously shown (*25–27*) to directly disrupt intestinal epithelial barrier function, resulting in increased monolayer permeability as observed in co-culture with CD4act in Figure 2C. An increase in FasL in the CD4act co-culture supernatants is consistent with the reduced viability of LGR5⁺ stem-cell observed in enteroids co-cultured with CD4act cells, as observed in Figure 1E.

**Figure. 3:**
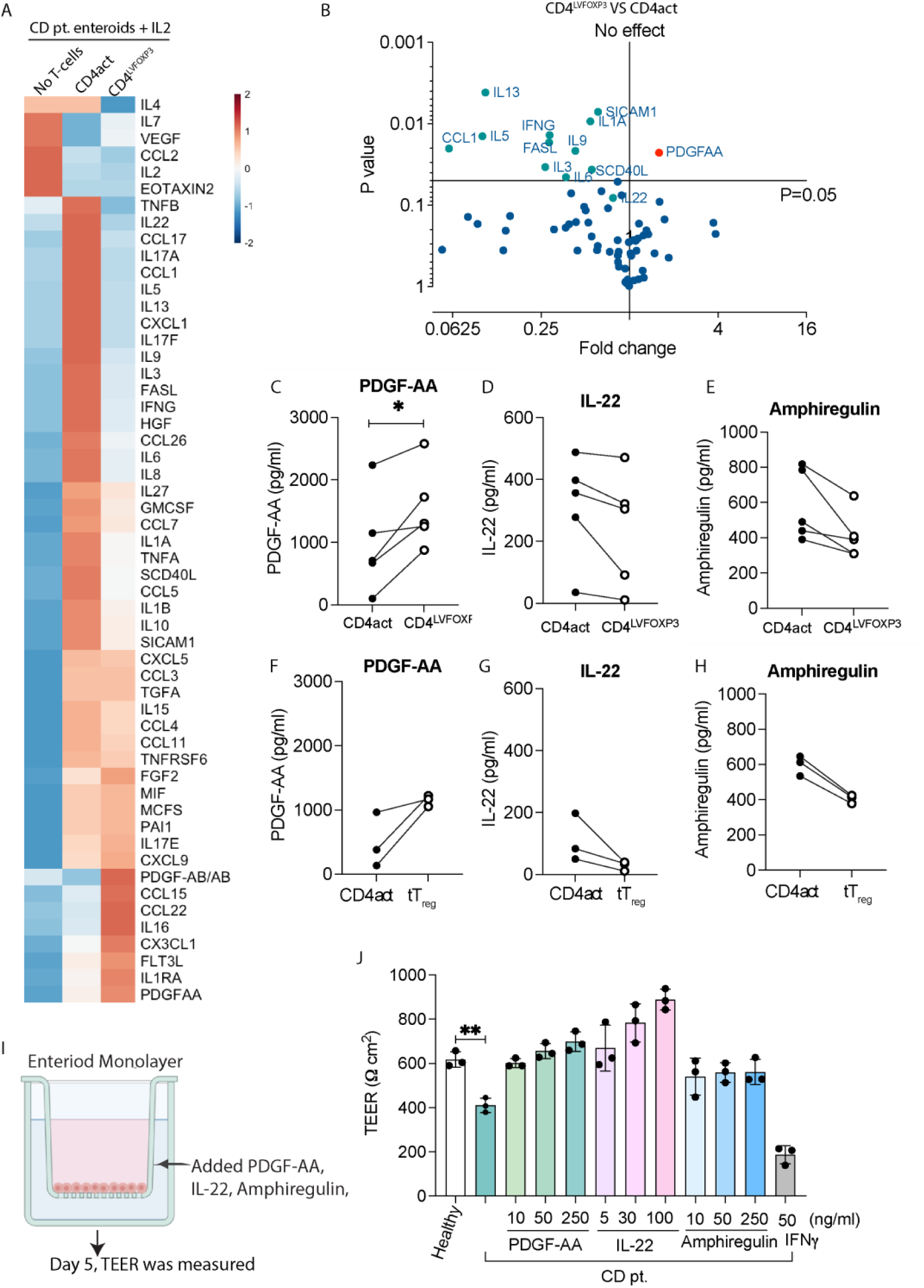
CD4^LVFOXP3^ and tT_reg_ cells reduce pro-inflammatory cytokines and enrich epithelial repair factors in the co-culture secretome. **(A)** Heatmap depicting the mean concentrations of the indicated cytokines and proteins in supernatant from enteroids monolayer only or CD4^LVFOXP3^ or CD4act co-culture. **(B)** Volcano plot showing the expression of the indicated proteins in the co-culture supernatant from enteroids cultured with CD4^LVFOXP3^ compared to CD4act. **(C-H)** Concentration (pg/ml) of PDGF-AA (C, F), IL-22 (D, G), and amphiregulin (E, H) in supernatant from enteroid monolayer and indicated cells co-cultures**. (I)** Schematic showing the experimental setup for determining the effect of PDGF-AA, IL-22, or amphiregulin on the TEER. **(J)** TEER measured in monolayer cultured without or with PDGF-AA, IL-22, Amphiregulin or IFNγ at indicated concentrations. Data plotted are from independent experiments performed with five CD patients (A-E) or three patients (F-J). * indicates statistically-significant difference (paired t-test, p > 0.05).

A consistent statistically significant (p=0.02) increase in the growth factor PDGF-AA was detected in CD4^LVFOXP3^ co-cultures (**Fig. 3B, 3C**). IL-22, a key cytokine that supports gut barrier integrity and epithelial repair (*11, 28*) and Amphiregulin, a tT_reg_ cell-secreted effector protein implicated in lung and thymic epithelial tissue repair (*29, 30*) were detectable in CD4^LVFOXP3^ and tT_reg_ co-culture supernatants but at lower levels than that of CD4act co-cultures (**Fig. 3D, E, G, H**). Similarly, supernatants from enteroid–tT_reg_ cells co-cultures exhibited significantly lower levels of inflammatory cytokines compared with those from CD4act cells from the same donor–enteroids co-cultures (**fig. S4C**). In contrast, IL-7, CCL22, and PDGF-AA levels were increased, although not significantly, in enteroid– tT_reg_ cells supernatants compared to those from CD4act co-culture (**fig. S4C, Fig. 3F**). Together, these data indicate that co-culture supernatants from both CD4^LVFOXP3^ and tT_reg_ cells showed reduced levels of pro-inflammatory cytokines while maintaining the presence of mediators associated with epithelial repair. This finding is consistent with the observation that in these co-cultures, TEER values were increased. In contrast, co-culture supernatant from CD4act cells showed significantly elevated levels of inflammatory cytokines, which correlate with TEER disruption, loss of polarity, and reduced viability of LGR5^+^ cells. To determine the sufficiency of CD4^LVFOXP3^ and tT_reg_ secreted proteins for bolstering epithelial barrier function, we tested the effect of recombinant PDGF-AA, IL-22, and amphiregulin on TEER values from CD enteroid-derived monolayer (**Fig. 3I**). The addition of each of these soluble factors increased TEER values relative to control, with PDGF-AA and IL-22 alone eliciting the largest effects (**Fig. 3J**); nonetheless, TEER values achieved by any of these single soluble factors remained lower than that observed in the presence of CD4^LVFOXP3^ (**Fig. 3J**). These results suggest that while PDGF-AA and IL-22 are sufficient to augment epithelial barrier function, additional mediators produced by CD4^LVFOXP3^ may act synergistically.

### PDGF-AA is essential for tT_reg_/CD4^LVFOXP3^-driven improvements in epithelial TEER and enteroid size

To test whether CD4^LVFOXP3^ or tT_reg_ derived PDGF-AA directly contributes to the enhancement of TEER and enteroids size, we first determined whether ISCs in our system are responsive to PDGF-AA via PDGFRα. Stimulation of enteroid with recombinant PDGF-AA selectively increased AKT phosphorylation at S473 without affecting ERK phosphorylation at T202/Y204, indicating the preferential activation of PDGFRα–AKT axis (**fig. S5A**). Next, we found that pharmacological inhibition of PDGFRα with avapritinib (*31*) abrogated the CD4^LVFOXP3^–induced increase in TEER in CD enteroid monolayers relative to vehicle control (**Fig**. **4A, B**). In this experiment, CD4^LVFOXP3^ co-culture enhanced pAKT(S473) protein levels, which were abolished upon PDGFRα inhibition (**Fig. 4B inset)**. Furthermore, antibody neutralization of PDGF-AA significantly reduced the CD4^LVFOXP3^-dependent TEER and pAKT(S473) compared with the isotype control **(Fig. 4C).** Importantly, anti–PDGF-AA had no effect on TEER in control epithelial monolayers alone but significantly reduced (p value=0.016) TEER in monolayers co-cultured with either CD4^LVFOXP3^ or tT_reg_ cells **(Fig. 4D)**, consistent with a paracrine T_reg_-to-ISCs PDGF-AA signaling. Undifferentiated enteroids co-cultured with tT_reg_ cells exhibited increased size, comparable to that observed in enteroids co-cultured with CD4^LVFOXP3^ cells in Figure 1I (**Fig. 4E)**. Blocking PDGF-AA resulted in significantly reduced (p value=0.0002) size of enteroids co-cultured with CD4^LVFOXP3^ or tT_reg_ relative to controls (**Fig. 4F**). These data indicate that PDGF-AA produced by CD4^LVFOXP3^ cells or tT_reg_ cells, signals via epithelial PDGFRα and is necessary to promote enteroid growth and epithelial barrier function.

**Figure 4:**
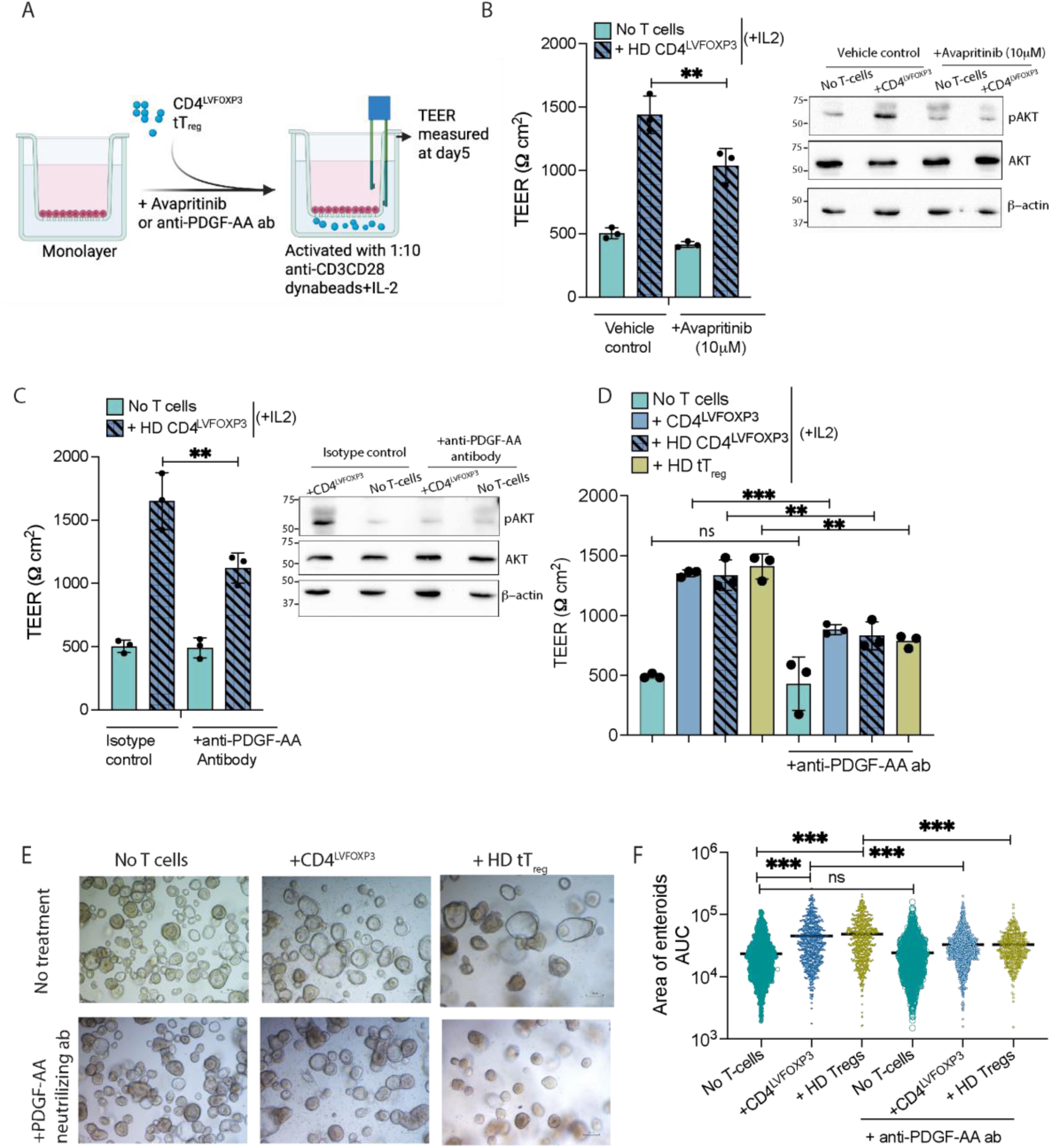
Inhibition of PDGF-AA signaling abrogates CD4^LVFOXP3^-mediated improvement of TEER and enteroid size. **(A)** Schematic showing the experimental setup to assess the inhibition of the PDGF-AA signalling. **(B, C)** TEER in monolayer co-cultured without or with allogenic CD4^LVFOXP3^ and treated with the 10μM Avapritinib (PDGFRα inhibitor) or vehicle control **(B)** or anti-PDGF-AA (0.9 μg/ml) or Isotype antibody **(C)**. Insets showing immunoblot probed for pAKT (S473), total AKT and β-actin in enteroids monolayer cultured alone or with CD4^LVFOXP3^ with indicated treatments. **(D)** TEER in monolayer co-cultured without or with autologous CD4^LVFOXP3^, allogenic CD4^LVFOXP3^ or tT_reg_ cells and treated with anti-PDGF-AA antibody (0.9 μg/ml). **(E, F)** Representative brightfield images of enteroids co-cultured without or with autologous CD4^LVFOXP3^ or allogenic CD4^LVFOXP3^ or tT_reg_ cells treated with the anti-PDGF-AA antibody. (E) and quantified for the size (F). Data plotted are from independent experiments performed with three patients. Statistical analyses were performed using paired t-test (C-F, J). ** and *** denote p < 0.01 and p < 0.001, respectively; ns denotes p > 0.05.

### CD4^LVFOXP3^ and tT_reg_ cells inhibit CD4act-induced damage independent of PDGF-AA

Previous experiments indicated that PDGF-AA contributes to the CD4^LVFOXP3^-mediated increases in enteroid size and TEER value, but its role in the CD4^LVFOXP3^ suppressive function was not assessed. To determine whether PDGF-AA is required for the immunosuppressive activity excreted by T_reg_, we established a three-component co-culture system in which enteroids were co-cultured with CD4act T cells together with either CD4^LVFOXP3^ cells or tT_reg_ cells (**Fig. 5A**). In enteroid–T cell co-cultures from 4 independent CD donors, CD4act reduced the frequency of LGR5^+^ cells as previously shown in Figure 1H (**Fig. 5B**), whereas adding equal amounts of CD4^LVFOXP3^ or tT_reg_ cells to CD4act, prevented the reduction in LGR5^+^ cells (**Fig. 5B**). In these conditions, PDGF-AA blockade does not affect reduction of LGR5^+^ cells by CD4act cells and also does not attenuate the protection conferred by CD4^LVFOXP3^ or tT_reg_ cells (**Fig. 5B**), indicating that PDGF-AA is dispensable for their ability to block CD4act mediated damage.

**Figure 5.**
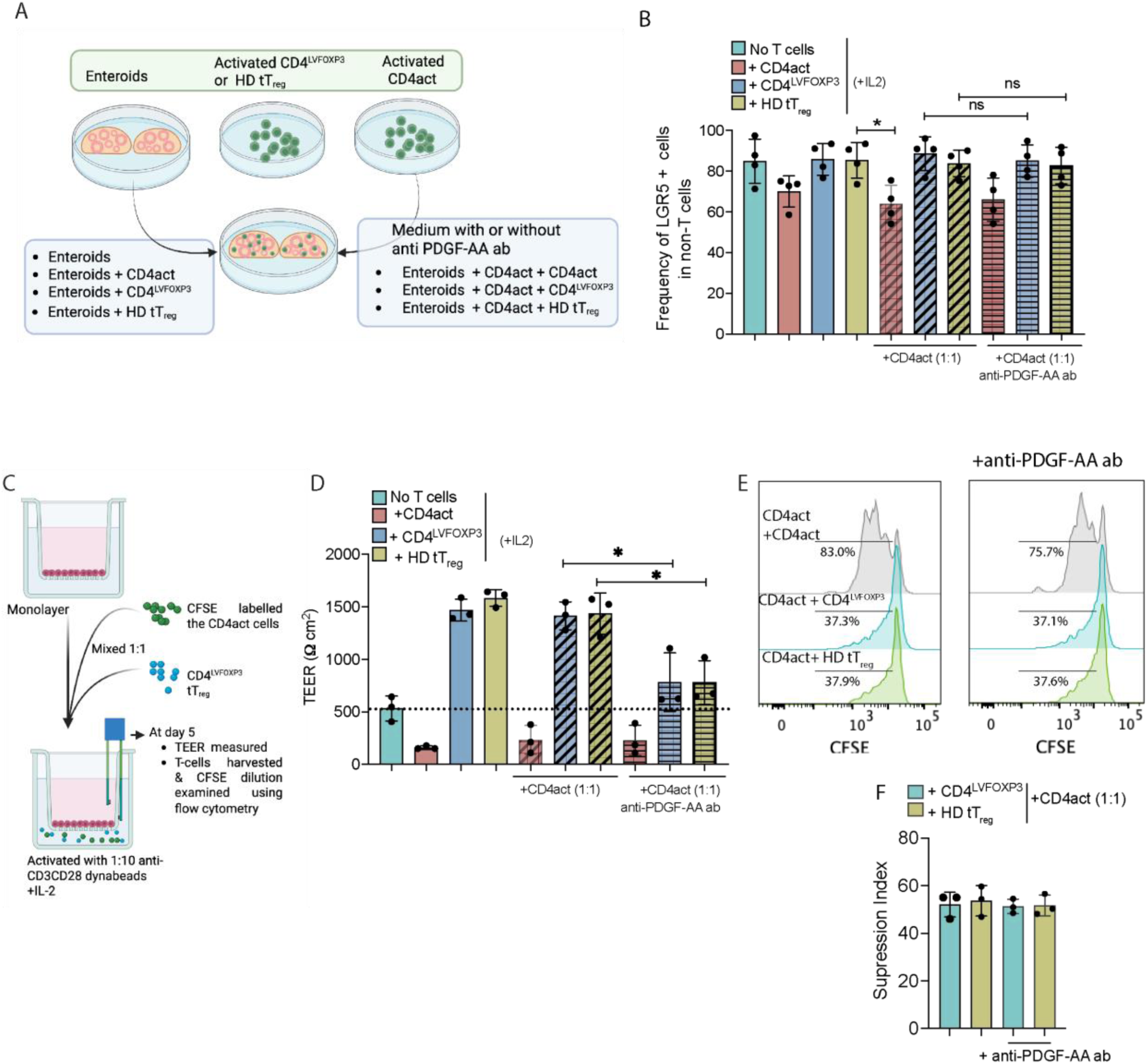
PDGF-AA signaling is dispensable for CD4^LVFOXP3^ or tT_reg-_mediated suppression of the CD4act. **(A)** Schematic showing the experimental setup of the co-culture of enteroid with CD4act or along with CD4^LVFOXP3^ or tT_reg_ cells. **(B)** Frequency of the LGR5^+^ cells in non-T cells in enteroids co-cultured as indicated in the schematic. **(C)** Schematic showing the experimental setup of the co-culture of enteroid with CD4act labelled with CFSE or along with autologous CD4^LVFOXP3^ cells or allogenic HD tT_reg_ cells. **(D-F)** TEER measured in monolayers from CD patients co-culture with the indicated cell types (D). Representative histograms showing CFSE dilution in activated CD4⁺ T cells cultured under the indicated conditions, in the presence or absence of anti–PDGF-AA antibody (E). Quantification of the suppression index mediated by autologous CD4^LVFOXP3^ cells or allogenic HD tT_reg_ cells (F). Data plotted are from independent experiments performed with four (B) or three (D-F) CD patients. Statistical analyses were performed using a paired t-test (C-F, J). * denote p < 0.01; ns non-significant denotes p > 0.05.

To concurrently assess the requirement of PDGF-AA in immunoregulatory activity and ISCs support within a single assay, we co-cultured enteroid monolayers with CD4act alone labeled with Carboxyfluorescein succinimidyl ester (CFSE) dye or together with equal amounts of CD4^LVFOXP3^ or tT_reg_ cells. Epithelial barrier integrity was measured by TEER as readout of CD4^LVFOXP3^ repair capacity, and inhibition of proliferation of activated CD4^+^ T cells—as a proxy for their suppression—was quantified by CFSE dilution (**Fig. 5C**). Co-culture with CD4^LVFOXP3^ cells or tT_reg_ cells prevented the CD4ac mediated decrease in TEER and even elevated TEER values above the levels in monolayers cultured alone (**Fig. 5D**), indicating that these regulatory populations both blocked CD4act-mediated damage and actively enhanced epithelial barrier function. In the same experiment, proliferation of CD4act was inhibited by the addition of the CD4^LVFOXP3^ cells or tT_reg_ cells (**Fig. 5E**). PDGF-AA neutralization had no effect on the CD4act-mediated decrease in TEER values but significantly reduced TEER in monolayers co-cultured with CD4act plus CD4^LVFOXP3^ cells or tT_reg_ cells, although TEER remained higher than in monolayers cultured without T cells (**Fig. 5D**). PDGF-AA blockade did not impair CD4^LVFOXP3^ or tT_reg_-mediated inhibition of CD4act proliferation (**Fig.5E, F**). Together, these results indicate that PDGF-AA is required for the CD4^LVFOXP3^/tT_reg_-mediated enhancement of epithelial barrier function but is dispensable for their suppression of CD4act.

### PDGFRα-PDGFA is expressed in ISCs and T_reg_ cells, respectively in CD patients

We asked whether PDGFRα is enriched in LGR5^+^ ISCs and if intestinal FOXP3^+^ T_reg_ upregulate PDGF-AA in CD patients. We showed that LGR5^+^ ISCs, which are enriched in undifferentiated enteroids, express the PDGFRα receptor at comparable levels between CD and HD enteroids, as revealed by quantitative immunoblot analysis (**Fig. 6A**). Next, to assess whether PDGFRα expression was restricted to the LGR5^+^ cell compartment, enteroids were differentiated to evaluate different cell types as in Figure 1H and PDGFRα protein levels was found to be reduced in differentiated versus undifferentiated enteroids from CD (**Fig. 6B**), consistent with its preferential expression in the LGR5^+^ stem-cell fraction. To confirm this finding, we examined PDGFRα expression in differentiated enteroids, which contained both LGR5^−^ cells (putative differentiated epithelial populations) and LGR5^+^ cells. Among LGR5^−^ cells, fewer than 40% were PDGFRα^+^, whereas more than 80% of LGR5^+^ cells were PDGFRα^+^, indicating that PDGFRα and LGR5 are co-enriched in the same cell population (**Fig. 6C**).

**Figure. 6.**
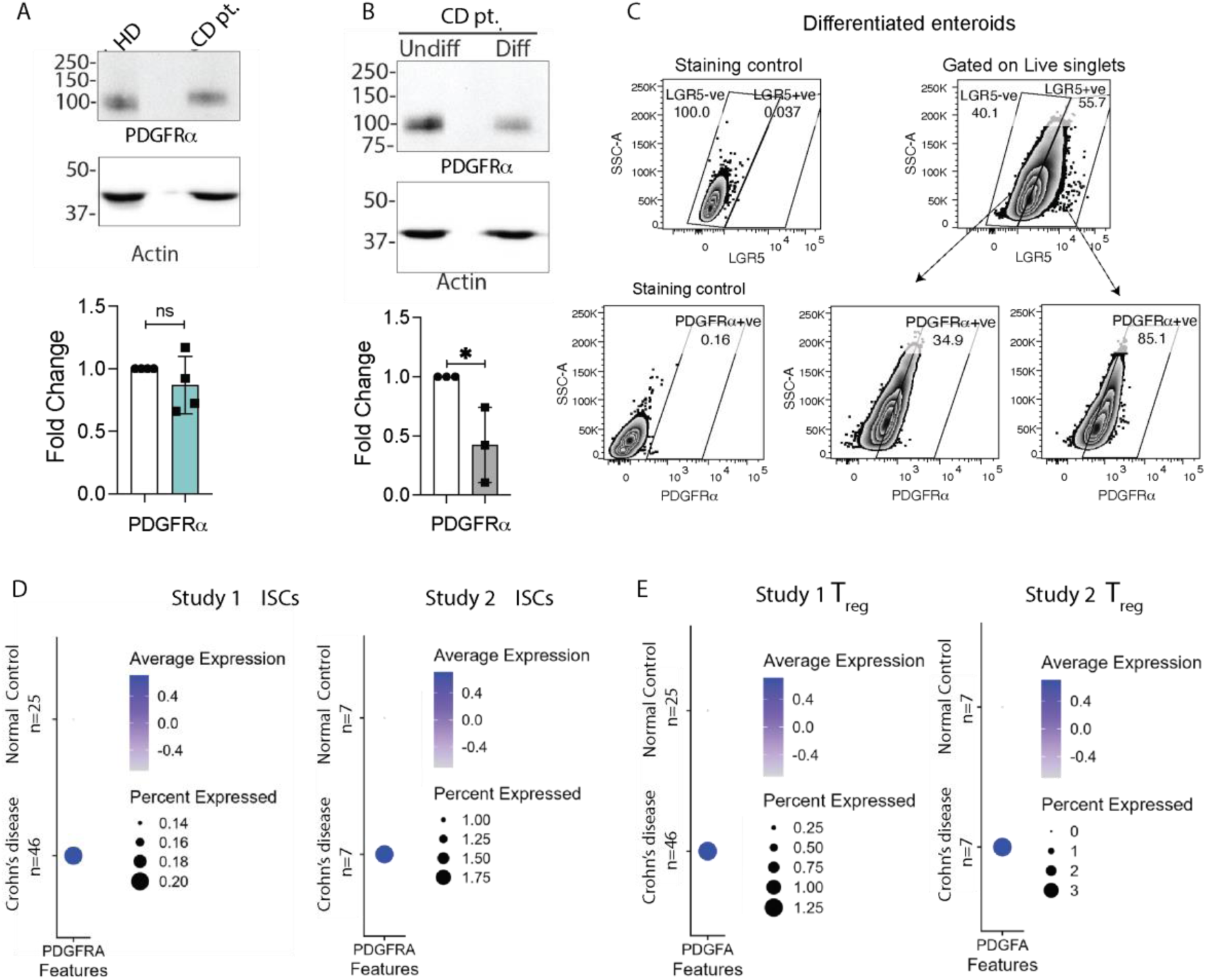
PDGFRα-PDGFA is expressed in ISCs and T_reg_ cells, respectively. **(A)** Representative immunoblot of cell lysates from enteroids from CD patients or HD probed for PDGFRα. Inset showing fold change in the expression levels of the PDGFRα in CD patients compared to HD. **(B)** Representative immunoblot of cell lysates from enteroids from CD patients differentiated for 5 days and probed for PDGFRα. Inset showing fold change in the expression levels of the PDGFRα in differentiated as compared to undifferentiated enteroids. **(C)** Representative flow cytometry plots showing PDGFRα+ cells within LGR5^−^ and LGR5^+^ populations. **(D)** Expression of PDGFRA gene in stem cells analyzed by single-cell sequencing from Study 1 and Study 2. **(E)** Expression of PDGFA gene in T_reg_ cells analyzed by single-cell sequencing from Study 1 and Study 2. Data plotted are from independent experiments performed with three CD patients or HD. Statistical analyses were performed using an unpaired t-test. * denote p < 0.01; ns denotes non-significant, p > 0.05.

We then re-analyzed published single-cell RNA-sequencing datasets from adult ileum study 1 ((*32*) n=46 CD, n=25 Normal control) and pediatric ileum study 2 ((*33*) ; n=7 CD, n=7 Normal control) to assess expression of PDGFRA gene and its ligand PDGFA across intestinal epithelial and immune compartments. Within the epithelial compartment, PDGFRA transcripts were detected in only a small fraction of cells, with expression observed not only in ISCs but also in enterocytes and enteroendocrine cells (**fig. S6A, B).** Across both cohorts, PDGFRA transcripts were enriched in CD-derived ISCs clusters relative to normal controls (**Fig. 6D**). In the immune compartment, PDGFA expression was highest in macrophages and monocytes in the adult dataset, with detectable expression in T_reg_ cells (**fig. S6C**), whereas in the pediatric cohort, PDGFA expression was highest in dendritic cells and was also present in T_reg_ cells (**fig. S6D**). Notably, PDGFA expression was increased in FOXP3^+^ T_reg_ clusters from CD compared to HD (**Fig. 6E**). Taken together, these data suggest that PDGFRα is expressed by ISCs and is maintained in CD-derived ISCs, whereas PDGFA is increased in FOXP3^+^ T_reg_ cells in CD, supporting the existence of a disease-associated PDGF-A–PDGFRα epithelial–immune signaling axis in the inflamed ileum.

## DISCUSSION

In the present study, we demonstrate that, beyond their primary immunomodulatory role, engineered T_reg_, CD4^LVFOXP3,^ similarly to tT_reg_ cells, exhibit a distinct and previously unreported therapeutic potential for directly enhancing intestinal epithelial growth and barrier function. Using patient-derived intestinal enteroid-T-cells co-culture, we found that the presence of the T_reg_ cells maintained the LGR5^+^ ISC pool and enhanced enteroid growth, without affecting their differentiation capacity. Given that ISC stemness and renewal are impaired in IBD, including CD (*34–36*), our data support a role for T_reg_ cells in ISC self-renewal and, consequently, intestinal epithelial regeneration. Our results show that CD–associated epithelial barrier dysfunction was preserved in enteroid monolayers, as reflected by reduced TEER and altered apical–basal polarity of ISC, indicating an epithelial-intrinsic defect that persists *ex vivo* and provides a suitable experimental model to test the impact of the T_reg_ cells. Indeed, co-culture with CD4^LVFOXP3^ cells or tT_reg_ cells rescued barrier function and restored polarity, suggesting that T_reg_ cells strengthen epithelial junctions. CD4^LVFOXP3^ and tT_reg_ had comparable effects on restoring epithelial barrier function, suggesting a shared FOXP3-dependent mechanism driving epithelial repair. Because our enteroid–T cell co-cultures exclude stromal and myeloid compartments, the rescue observed here reflects epithelial-intrinsic responsiveness to T_reg_-derived factors under defined conditions. Collectively, these data provide evidence for a functional crosstalk between T_reg_ cells and the human ISCs compartment, and for T_reg_-mediated restoration of epithelial barrier function—outcomes not captured in earlier studies focused largely on indirect, immune-centered mechanisms of mucosal healing (*9, 10, 37*).

Mechanistically, the effect of T_reg_ on ISCs was associated with a favorable secretome profile: low levels of pro-inflammatory cytokines, alongside the presence of tissue-remodeling growth factors such as PDGF-AA, IL-22, and amphiregulin. In our system, recombinant IL-22, PDGF-AA and amphiregulin directly enhanced barrier function, with IL-22 and PDGF-AA showing the strongest effects. IL-22 has been shown to support epithelial regeneration and Paneth-cell programs essential for ISC maintenance (*38, 39*). T_reg_-derived amphiregulin—although implicated in epithelial repair in the lung and thymus (*29, 30*)—increased TEER, with an effect size smaller than IL-22 or PDGF-AA. Thus, amphiregulin-EGFR signaling alone may be insufficient to strengthen epithelial barrier integrity in the intestinal context. Together, the data we describe provide evidence for a T_reg_-driven, multi-factor regenerative program that supports ISC maintenance, at least in part, through engagement of the PDGF-AA–PDGFRα signaling axis. Our re-analysis of single-cell RNA-seq datasets supports this mechanism in human tissue. PDGFRA transcripts were detected in ISCs, whereas PDGFA transcripts were detected in intestinal T_reg_ cells. Expression of both genes was increased in CD as compared with HD, suggesting a role for PDGF-AA-PDGFRα signaling axis in epithelial repair programs in inflamed tissue. Tissue-resident T_reg_ cells in other barrier organs (colon, lung, skin) also expressed PDGFA gene, suggesting a conserved program whereby T_reg_-derived PDGF-AA provides trophic support to epithelial progenitors across barrier sites (*40*). Our data define a FOXP3-dependent, Treg-driven PDGF-AA–PDGFRα axis that directly supports the ISC niche and epithelial barrier repair, extending beyond previously described T_reg_-mediated amphiregulin–EGFR, and IL-10 pathways, while complementing epithelial-intrinsic IL-22 signaling. The PDGF-AA–PDGFRα pathway regulates gut epithelial homeostasis through PDGFRα+ pericryptal stromal cells (*41, 42*). In this context, PDGF-AA secreted by T_reg_ cells could act on pericryptal PDGFRα+ stromal cells, which are a core component of the ISC niche and supply essential Wnt agonists (Wnts, R-spondins), BMP antagonists, and other trophic factors (*41–43*). Thus, T_reg_-derived PDGF-AA may help sustain niche architecture and indirectly stabilize the signals required for ISC fitness. Importantly, PDGF-AA was dispensable for T_reg_-mediated suppression of CD4^+^ T cells proliferation.

In contrast to T_reg_, conventional autologous activated CD4^+^ T cells from CD peripheral blood damaged ISCs as shown by reduced viability and altered differentiation capacity. Although these cells were blood-derived, their pro-apoptotic effect resembles that reported for mucosal effector T-cells in analogous co-culture systems (*44*). Additionally, the activated/effector CD4^+^ T-cells from CD abrogated apical–basal polarity and reduced TEER below baseline, supporting the notion that effector T-cells directly perturb epithelial homeostasis. Their secretome profiling revealed high concentrations of barrier-disruptive inflammatory mediators, including IFN-γ and IL-13, together with FASL, providing a mechanistic link between effector T cell activation and epithelial injury (*25–27*). Together, we propose an opposing T cell–epithelial interactions with T_reg_ cells maintaining ISCs and enhancing epithelial barrier function, whereas activated CD4^+^ T cells are damaging ISCs and impairing barrier integrity.

The observation that blocking PDGF-AA–PDGFRα signaling did not completely abrogate the T_reg_-mediated improvement in barrier function indicates the involvement of synergistic factors, including IL-22 and amphiregulin. Defining the necessity and sufficiency of IL-22, amphiregulin–EGFR signaling, and additional mediators improving the barrier function will be an important future direction. Our ISC–T_reg_ co-culture system delineated epithelial-intrinsic responses to tT_reg_-derived signals under defined conditions. However, this approach is inherently limited by the absence of key *in vivo* cellular components that shape crypt homeostasis and barrier repair, including PDGFRα⁺ mesenchymal/stromal populations, myeloid and other lymphoid lineages. Because stromal and myeloid compartments provide essential niche cues, future studies will require integrated epithelial–stromal and immune-inclusive models to define how T_reg_-derived signals converge to promote epithelial repair.

Overall, the findings described here have translational relevance for CD, where current therapies—including broadly acting agents and targeted biologics—inhibit immune-cell migration or counteract inflammatory cytokines (*45*). Although these treatments alleviate the damage, they do not directly promote epithelial healing or result in long-term remission. In this context, T_reg_ cells may be able to play a dual role in re-establishing the immune homeostasis and favoring repair. T_reg_ cells are, in principle, naturally well-suited to re-establish mucosal homeostasis and promote repair; however, inflammatory milieus can compromise gut-resident T_reg_ stability and function (*46*). Functional T_reg_ stability indeed remains a challenge, also for T_reg_ therapies in which the cell product is obtained upon *in vitro* expansion. By contrast, engineered CD4^LVFOXP3^ cells, which exhibit a defined and extensively characterized regulatory phenotype, may provide a more robust strategy by coupling suppression of inflammation with active promotion of epithelial restitution in CD and other immune-mediated enteropathies. CD4^LVFOXP3^ cells have been shown to control immune pathology in humanized mouse models of Graft-versus-host disease (GVHD) and IPEX, and to migrate to intestinal tissue *in vivo* (*12, 13*). Building on these observations, the present pre-clinical study establishes that CD4^LVFOXP3^ cells can rescue barrier function and protect the intestinal stem-cell compartment from damage driven by activated CD4^+^ T cells.

## MATERIALS AND METHODS

### Study Design

This study was designed to investigate the interaction between T_reg_ and human ISCs. Primary human intestinal enteroids were established from the CD (n=9) patients or HD (n=8) ileal tissue. T_reg_ cells, both engineered CD4^LVFOXP3^ or tT_reg_ were generated from CD (n=5) or HD (n=3) peripheral blood and co-cultured with autologous or allogeneic enteroids. Sample size was not determined by an a priori power analysis. This study was designed to be exploratory in nature, and the sample size was selected based on feasibility (availability of paired ileal tissue and peripheral blood specimens) and on achieving stable estimates of the primary outcome. ISCs maintenance, cell viability, epithelial polarity, and barrier integrity were evaluated as primary outcomes (n of samples per assay is indicated in figure legends). Based on the measurement of soluble factors in enteroid-T cells co-culture supernatant, studies were performed to delineate tT_reg_-derived mechanisms governing ISCs support, with particular emphasis on PDGF-AA/PDGFRα signaling regulation of epithelial integrity. Experiments were performed using biological replicates refer to independent human subject. Technical replicates refer to parallel wells/enteroids (and imaging fields, where applicable) measured within the same subject per condition.

Allocation of samples to conditions, assays, and processing order was randomized. For co-culture experiment, only patient’s enteroid and autologous CD4^LVFOXP3^ were included if both ileal tissue and peripheral blood were available. For allogenic co-culture, HD CD4^LVFOXP3^ and tT_reg_ were used CD patient’s enteroids. Outlier handling was not used to remove data points unless there was a technical assay failure.

Pediatric participants were enrolled through the Center for Pediatric IBD & Celiac Disease Biorepository at Lucile Packard Children’s Hospital, Stanford (Stanford, CA, USA). Participant inclusion/exclusion criteria were defined prospectively by the biorepository protocol. Study procedures for recruitment, endoscopic biopsy and blood collection were approved by the Stanford University Institutional Review Board (IRB: 63678); written informed consent was obtained from parents or legal guardians. Biopsies and/or blood were collected during routine, clinically indicated endoscopy as part of standard patient care. During the procedure, four to eight biopsies (2–5 mm^3^ each) of terminal ileum and/ or ∼9mL of peripheral blood in EDTA-coated tubes were collected. De-identified ileal tissues (surgically-resected specimens) from patients with CD and non-IBD controls were obtained from the pediatric surgery service under a consent waiver. Biopsies and tissues were placed in ice-cold DMEM/F12 supplemented with Normocin (50 µg/mL) and transported on ice to the laboratory for immediate enteroid derivation. Donor demographics are provided in Supplementary Table 1.

### Enteroid generation and culture

Enteroids were generated as described previously (*47*). Briefly, two biopsies (2–5 mm^3^) of ileum tissue were incubated in 2 mM EDTA in 1X PBS for 30 minutes at 4°C with rotation, then transferred to 2% sorbitol buffer (2% Sorbitol, 1% Sucrose, 1% BSA and 50 μg/mL Normocin in 1X PBS) in a petri dish. Crypts were mechanically dissociated using forceps under a dissecting microscope (Olympus SZ51/60) and were washed with wash medium (OWM) containing DMEM/F12, 10% FBS, 1X Penstep, 2mM GlutaMAX, and 10% FBS and mixed with ice-cold Matrigel (1:2). 20 mL droplets of the crypt-Matrigel suspension were plated into 12-well plates and incubated at 37°C for 20 minutes to polymerize. 1 mL of OGM-CM (IntestiCult™ Intestinal Organoid Culture Media (OGM) containing 10 µM SB431542, 10 µM Y27632 and 50 µg/mL Normocin) was then added, and the media was changed every 2–3 days.

When enteroids reached approximately 100 µm in diameter, cultures were passaged. Enteroids were dislodged from plates with ice-cold 1X PBS, mechanically fragmented, transferred to 15 mL tubes, and pelleted at 400 × g for 5 minutes at 4 °C. The pellet was resuspended in 400 µL of prewarmed TrypLE Express, incubated at 37 °C for approximately 3 minutes, and triturated 10–12 times to dissociate. Cells were diluted in OWM, centrifuged at 300 × g for 5 minutes at 4 °C, gently resuspended, and plated as described above.

### Transepithelial electrical resistance (TEER) measurements

Enteroids were harvested in cold 1× PBS and were digested using TrypLE for 15 minutes to obtain single-cell suspension. Cells were washed with 5 mL OWM and centrifuged at 400 × g for 5 minutes. The resulting pellet was resuspended in 2 mL OGM medium and passed through a 70-µm cell strainer. A total of 3 × 10⁵ cells were seeded per well onto 0.4-µm pore size 24-well Transwell inserts in OGM supplemented with 50 µg/mL Normocin, 10 µM Y-27632, and 2 nM CHIR99021. After 24 hours, cultures were maintained in OGM containing 50 µg/mL Normocin and 10 µM Y-27632.

For differentiation, once confluent monolayers were established, the medium was replaced with ODM-CM (IntestiCult™ Intestinal Organoid Differentiation Media (ODM) supplemented with 50 µg/mL Normocin and 5 µM GSI-IX.

### Co-culture of enteroids and T cells

T cells (CD4^LVFOXP3^, CD4act, or tT_reg_) were thawed and rested overnight in X-VIVO 15 Medium supplemented with 5% human serum, 100 U/mL IL-2, and 1 ng/mL IL-15. After resting overnight, CD4^LVFOXP3^, CD4^NGFR^, or tT_reg_ cells were activated with Dynabead Human T cell Activator CD3/CD28 at a 1:10 cell number to bead ratio for 24 hours. Following activation, the Dynabeads were removed, and T cells were co-cultured with enteroids at a 1:100 ratio OGM-CM containing 100 U/mL IL-2 for 5 days. After 5 days, viability, the frequency of LGR5^+^ cells, and cell polarity were assessed in enteroids. For polarity analysis, enteroids were fixed in 4% paraformaldehyde and stained with Phalloidin-FITC for actin and an antibody to β-catenin as described above. For LGR5^+^ cells analysis, the single-cell suspension was immunostained for LGR5 and Live/Dead dye and analyzed using flow cytometry as described above.

Enteroid monolayers were prepared as previously described. Once the enteroid monolayers were established, overnight rested T cells were activated by adding anti-CD3 and CD28 Dynabeads at a 1:10 ratio in the bottom well of the transwell system. TEER readings were measured over 5 days to assess the impact of T cell activation on the integrity of the enteroid monolayer.

For TEER analysis in a differentiated monolayer, after the monolayer was formed, T cells were activated in the bottom well and cultured in ODM-CM and 100 U/mL IL-2. TEER was monitored for 4 days to evaluate the effects of T cell activation on the barrier function of the differentiated monolayer.

### Enteroid’ size measurement

Bright-field images were acquired using Nikon TS100 Inverted Phase Contrast Microscope at 10X, NA# 0.25. The images were processed using Cellpose-SAM for cellular segmentation with default settings and masks were generated for the identified organoids in the images (*48*). Subsequently, the organoid images and masks generated in Cellpose-SAM were imported into CellProfiler 4.2.8 (*49*). In CellProfiler, the Cellpose-SAM identified objects were filtered to exclude single cells and debris from the quantification. The area and other Zernike features of the filtered organoids were subsequently measured using the “Measure Object Size Shape” module in Cell Profiler and exported to spreadsheet.

### Re-analysis of Single-cell Sequencing

We utilized single-cell transcriptomes from the terminal ileum of childhood-onset CD and matched HD from two previously-published studies (*32, 33*). Processed single-cell expression matrices and metadata for human epithelial and immune cells for CD (n=46) samples and normal controls (n=24) were obtained from the Broad Single Cell Portal (https://singlecell.broadinstitute.org/single_cell, accession SCP1884) and for CD (n=7) samples and normal controls (n=7) from the Gut Cell Atlas (https://gutcellatlas.org). All cell type identities were maintained as reported in their studies of origin. Data processing and gene expression visualizations were performed using the R (v.4.3.3) package Seurat (v.5.4.0).

### Immunostaining and confocal Imaging

Enteroids were plated in Matrigel at a 1:1 ratio, plated on a confocal-grade chamber coverslip, and cultured for 24 hours in OGM-CM. After 24 hours, enteroids were fixed using 4% paraformaldehyde in 1X PBS for 30 minutes at room temperature in the dark. Enteroids were permeabilized with 0.02% Triton-X100 in 1X PBS for 15 minutes, followed by antigen retrieval using NH_3_Cl_2_ for 30 minutes at room temperature. Blocking was done using blocking buffer (0.01% TritonX-100, 1% BSA and 5% Normal goat serum) for 1 hour at room temperature. Primary antibody β-catenine (1:200) and FITC-phalloidin (1:300) were diluted in blocking buffer and incubated at 4 °C. The next day, enteroids were washed 3 times with 1X PBS and incubated with secondary antibody (1:500) in blocking buffer at room temperature for 1 hour in the dark. Then, enteroids were mounted using ProLong™ Glass Antifade Mountant with NucBlue™ Stain. Images were acquired SP8 Leica at 40X oil, NA # 1.40, at the Cell Sciences Imaging Facility, Stanford University.

### Luminex and ELISA analysis

Co-culture supernatants from enteroids monolayer alone or co-cultured with CD4^LVFOXP3^, tT_reg_, or CD4act were collected at the end of the experiment and centrifuged at 500 g for 5min at 4°C, aliquoted, and stored at −80°C until assay. Human 80-Plex Cytokine/Chemokine/Growth Factor Panels were analyzed using the Luminex xMAP^®^ platform at the Human Immune Monitoring Center (HIMC), Stanford University. Amphiregulin levels in supernatant were measured using the Human Amphiregulin DuoSet ELISA kit following the manufacturer’s instructions. Each biological sample was assayed in two technical replicates for analysis.

### Analysis of epithelial cell types in differentiated enteroids

Enteroids cultured in Matrigel were replated at a 1:2 ratio once they reached a diameter of 50–70 µm. Twenty-four hours after replating, OGM-CM was replaced with ODM-CM. Enteroids were cultured in ODM-CM for 5 days, with medium changes every other day.

Enteroids were harvested and washed three times with cold 1× PBS. Total RNA was isolated using the RNAqueous-Micro Total RNA Isolation Kit, and RNA concentration and purity were assessed by spectrophotometry. 1μg of RNA was treated with DNase I and reverse-transcribed into cDNA using the RevertAid RT Reverse Transcription Kit according to the manufacturer’s instructions. The resulting cDNA was diluted 1:5 and used for quantitative PCR with PowerUp SYBR Green Master Mix on a QuantStudio^TM^ Real-Time PCR System. Gene expression was quantified using the 2^−ΔΔCt^ method, with GAPDH as the reference gene.

GAPDH Fw: GACCTGCCGTCTAGAAAAACC Rv: GCTGTAGCCAAATTCGTTGTC CHGA Fw: AGAATTTACTGAAGGAGCTCCAAG Rv: TCCTCTCTTTTCTCCATAACATCC LGR5 Fw: TATGCCTTTGGAAACCTCTC Rv: CACCATTCAGAGTCAGTGTT SI Fw: CTGCATTTGAAAGAGGACAGC Rv: ACTCTGCTGTGGAAGTCCTGA MCU2Fw:AGGATCTGAAGAAGTGTGTCACTG Rv:TAATGGAACAGATGTTGAAGTGCT

### Statistical analysis

Data plotting and statistical analyses were performed using R (v2026.01.0+392) and GraphPad Prism (v7.0). Data are presented as mean ± s.d. Each dot represents an individual patient; the exact number of patients (n) is provided in the corresponding figure legends. For two-group comparisons, paired data (the same patient measured under different conditions) were analyzed using a two-tailed paired t-test. Unpaired data were analyzed using a two-tailed unpaired t-test with Welch’s correction when variances were unequal. All tests were two-sided, and P < 0.05 was considered statistically-significant.

## Supplementary Materials

Supplementary Methods

Figs. S1 to S5

Tables S1 to S2

## Acknowledgments

We would like to thank the patients and families who made this study possible by donating their blood and tissue biopsies. We thank Lee G. Fradkin for his insightful comments and thoughtful feedback on the manuscript. We thank the Stanford Institute for Stem Cell Biology and Regenerative Medicine FACS Core for the technical assistance. The Confocal Imaging was conducted at Cell Sciences Imaging Facility at Beckman Center with RRID SCR_017787; it was supported, in part, by Award Number 1S10OD010580-01A1 from the National Center for Research Resources (NCRR). Its contents are solely the responsibility of the authors and do not necessarily represent the official views of the NCRR or the National Institutes of Health.

## Author contributions

Conceptualization: N.S., R.B.; Methodology: N.S., B.J.O., P.G., J.A.L.; Investigation: N.S., B.J.O., N.B., A.R., P.G., J.A.L.; Visualization: N.S., J.A.L.; Funding acquisition: N.S., R.B., M.G.R.; Project administration: N.S., R.B.; Supervision: R.B., M.J.R.; Writing – original draft: N.S., R.B; Writing – review & editing: N.S., R.B., M.J.R., M.G.R., J.D.; B.J.O., A.R., P.G., J.A.L.

## Data and materials availability

All correspondence and requests for materials or coding scripts should be addressed to the corresponding author, R.B. All other data supporting the findings of this study are available within the paper and its Supplementary Materials.

## Supplementary Materials

**Suppl Fig 1:**
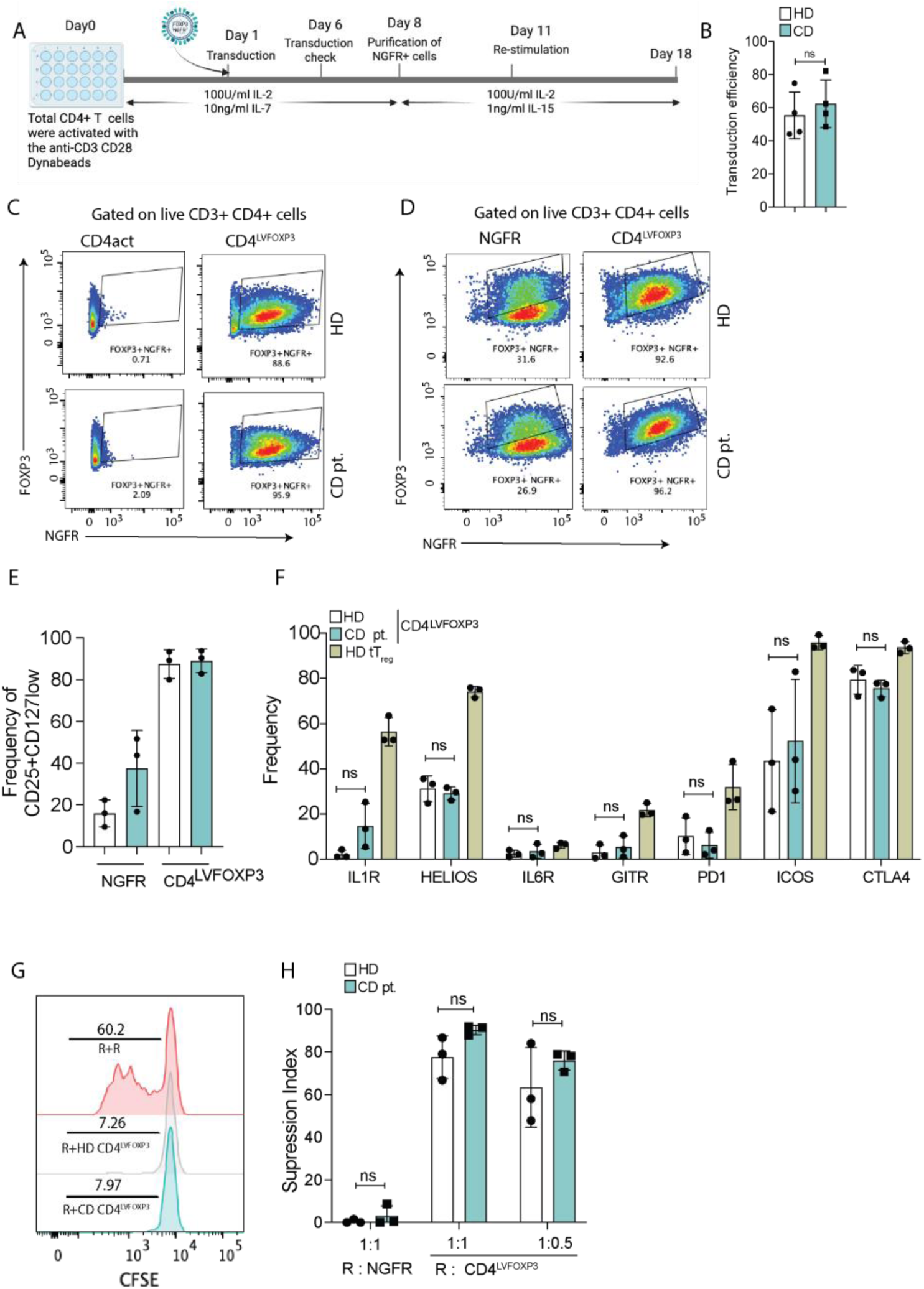
CD4^LVFOXP3^ cells generated from the CD patients and HD show a similar phenotype. **(A)** Schematic showing the protocol and key steps involved in the generation of CD4^LVFOXP3^ cells. **(B)** Frequency of NGFR+ cells before purification, indicating the transduction of the lentivirus in CD4 cells isolated from CD or HD PBMCs. **(C, D)** Representative flow plots showing the expression of FOXP3 and NGFR in CD4^LVOFPX3^ or Control CD4act cells (C) or CD4^NGFR^ cells (D) generated from CD patients and HD. **(E)** Frequency of CD25+ and CD127^low^ cells in CD4^LVOFPX3^ or CD4^NGFR^ cells from CD patients or HD. **(F)** Frequency of cells positive for the indicated protein in CD4^LVFOXP3^ cells generated from CD patients and HD and *in vitro* activated tT_reg_ from the same HD. **(G)** Representative Histogram plot showing CFSE dilution of responder (R) cells cultured with 1:1 R or CD4^LVOFPX3^ cells generated from CD patients or HD. **(H)** Suppression index of CD4^LVFOXP3^ or CD4^NGFR^ cells generated from CD patients and HD against Responder (R) cells. The data plotted are from experiments performed with the three CD patients or HD controls. ns indicates no statistically significant difference (paired t-test, p > 0.05).

**Suppl Fig. 2:**
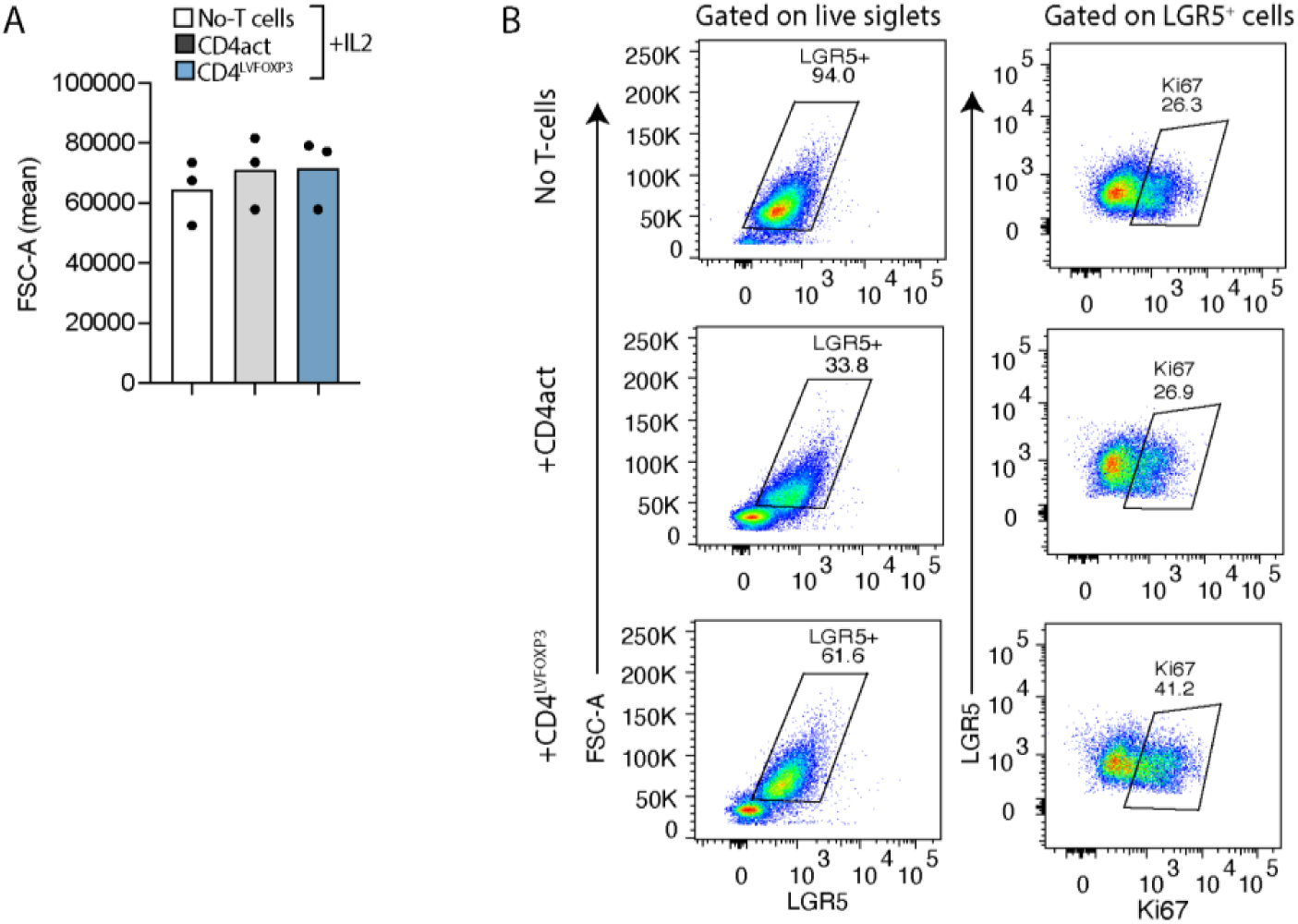
CD4^LVFOXP3^ increases the proliferation of the LGR5^+^ cells. (A-B) Forward scatter (FSC) mean (A) and Ki67+ (B) in LGF5^+^ cells in enteroids culture without T cells, CD4act or CD4^LVFOXP3^ on OGM+IL2 for 5 days.

**Suppl Fig. 3:**
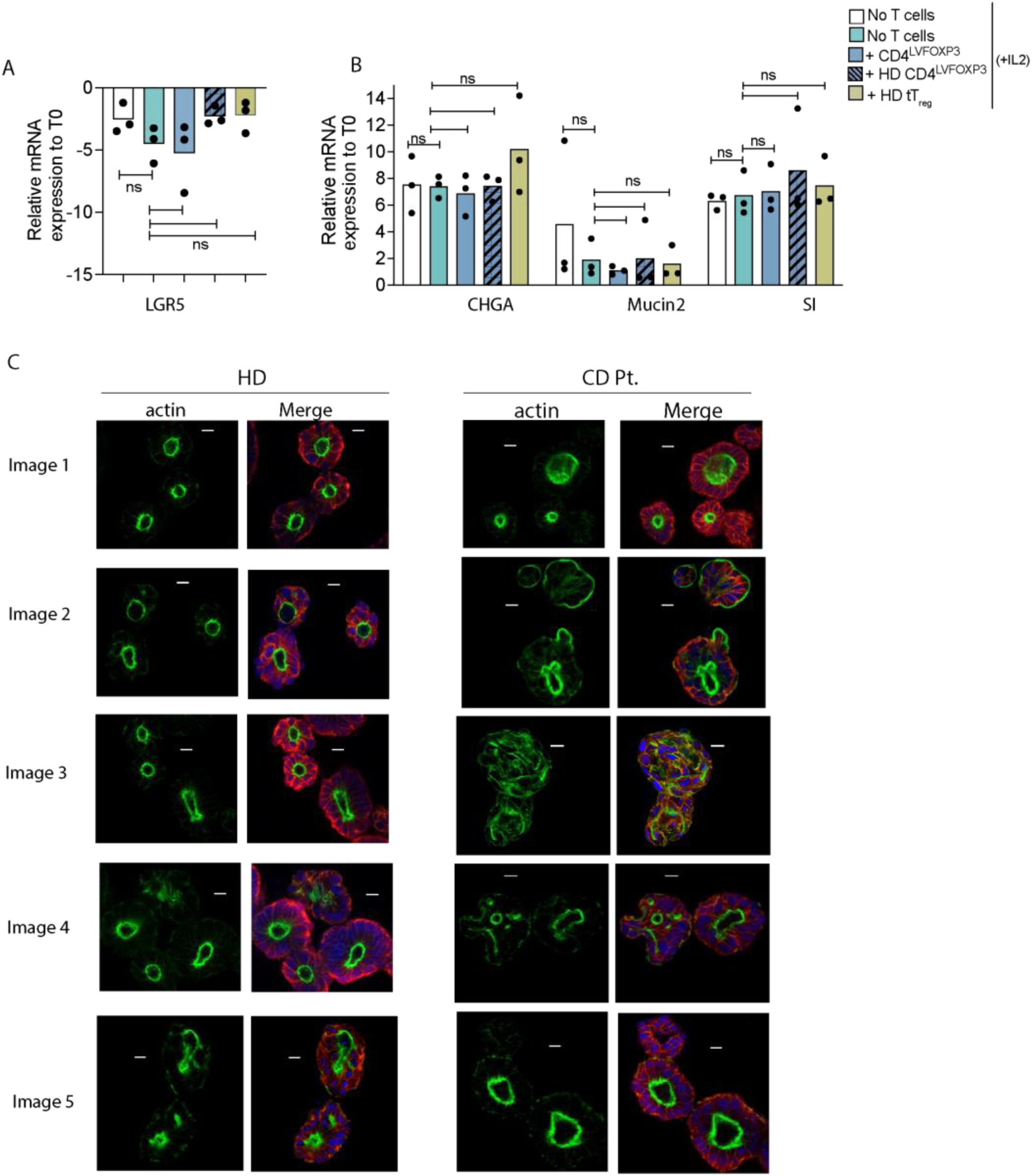
CD4^LVFOXP3^ and tT_reg_ do not affect enteroid monolayer differentiation. **(A, B)** Relative expression of LGR5 (A) and differentiation markers Chromogranin A (CHGA), Mucin 2 and Sucrose-isomaltase (SI) (B) in enteroids monolayer cultured alone or with indicated autologous and allogenic T cells in ODM-CM. **(C)** Representative confocal images of enteroids from CD patients or HD immunostained with Phalloidin actin (Alexa 488), anti-β-catenin antibody, and DAPI. Scale bar: 10μm. Data plotted are from independent experiments performed with CD (n=3) patients or HD (n=3) (A, B). ns indicates no statistically significant difference (paired t-test, p > 0.05).

**Suppl Fig. 4:**
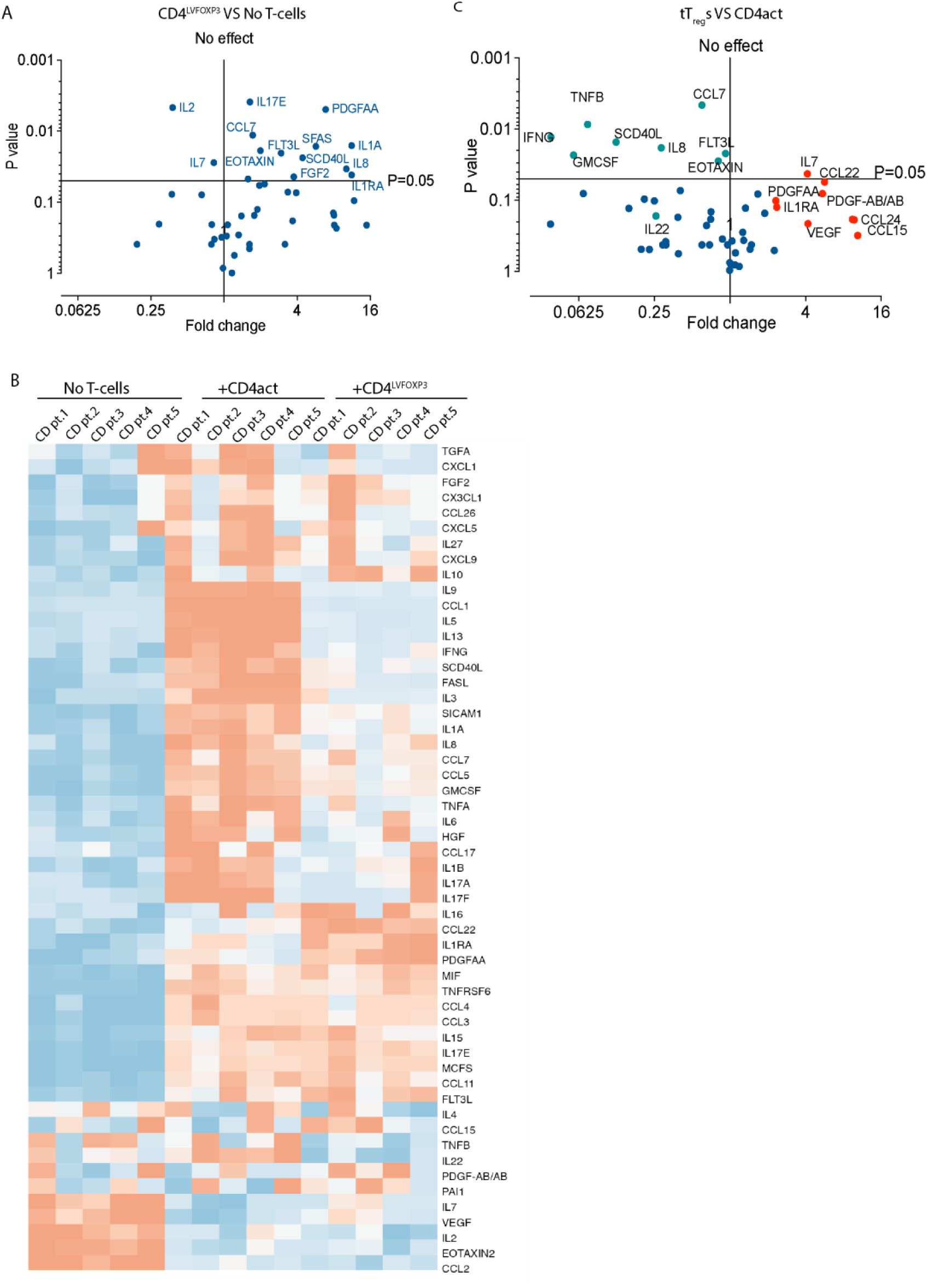
CD4^LVFOXP3^: enteroids monolayer co-culture supernatant shows reduced levels of pro-inflammatory cytokines. **(A)** Volcano plot showing the expression of the indicated proteins in the co-culture supernatant from enteroids monolayer co-cultured with CD4^LVFOXP3^ compared to monolayer only culture. **(B)** Heatmap showing the relative expression of cytokines and growth factors in supernatant from enteroid monolayer cultured without or with CD4^LVFOXP3^ or CD4act cells. (**C)** Volcano plot showing the expression of the indicated proteins in the co-culture supernatant from enteroids co-cultured with tT_reg_ compared CD4act cells. Data plotted are from independent experiments performed with five CD patients (A, B) or three patients (C).

**Suppl Fig 5:**
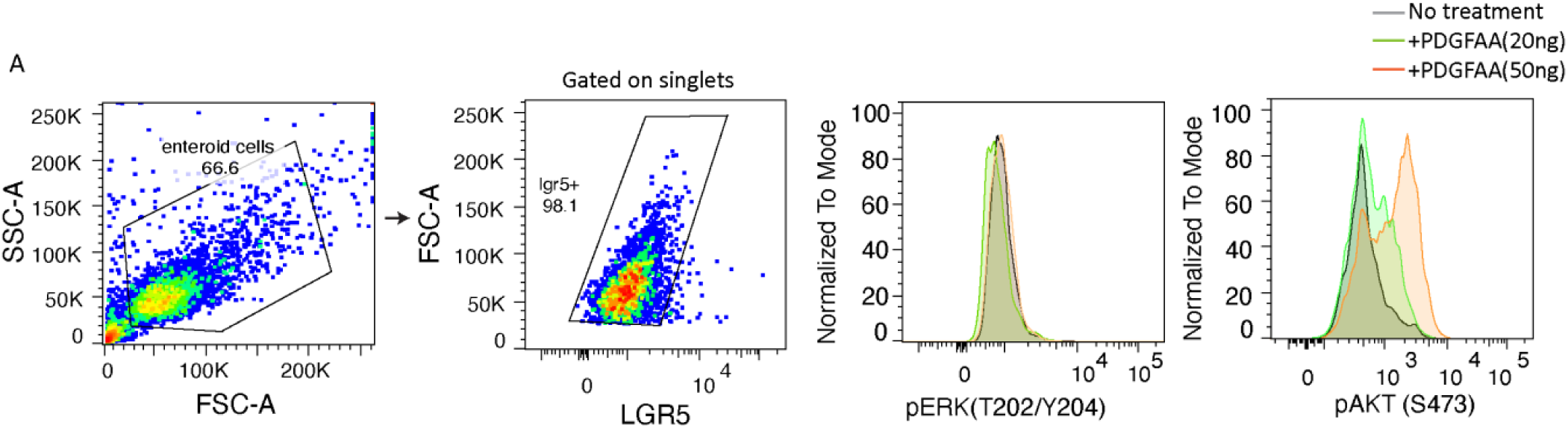
Enteroid cells respond to PDGF-AA. **(A)** Representative flow plot showing the expression of pERK (T202/Y204) and pAKT (S473) in enteroids cultured with PDGF-AA at the indicated concentration for 15 minutes. Data is representative of independent experiments performed with three CD patients-derived enteroids.

**Suppl Fig 6:**
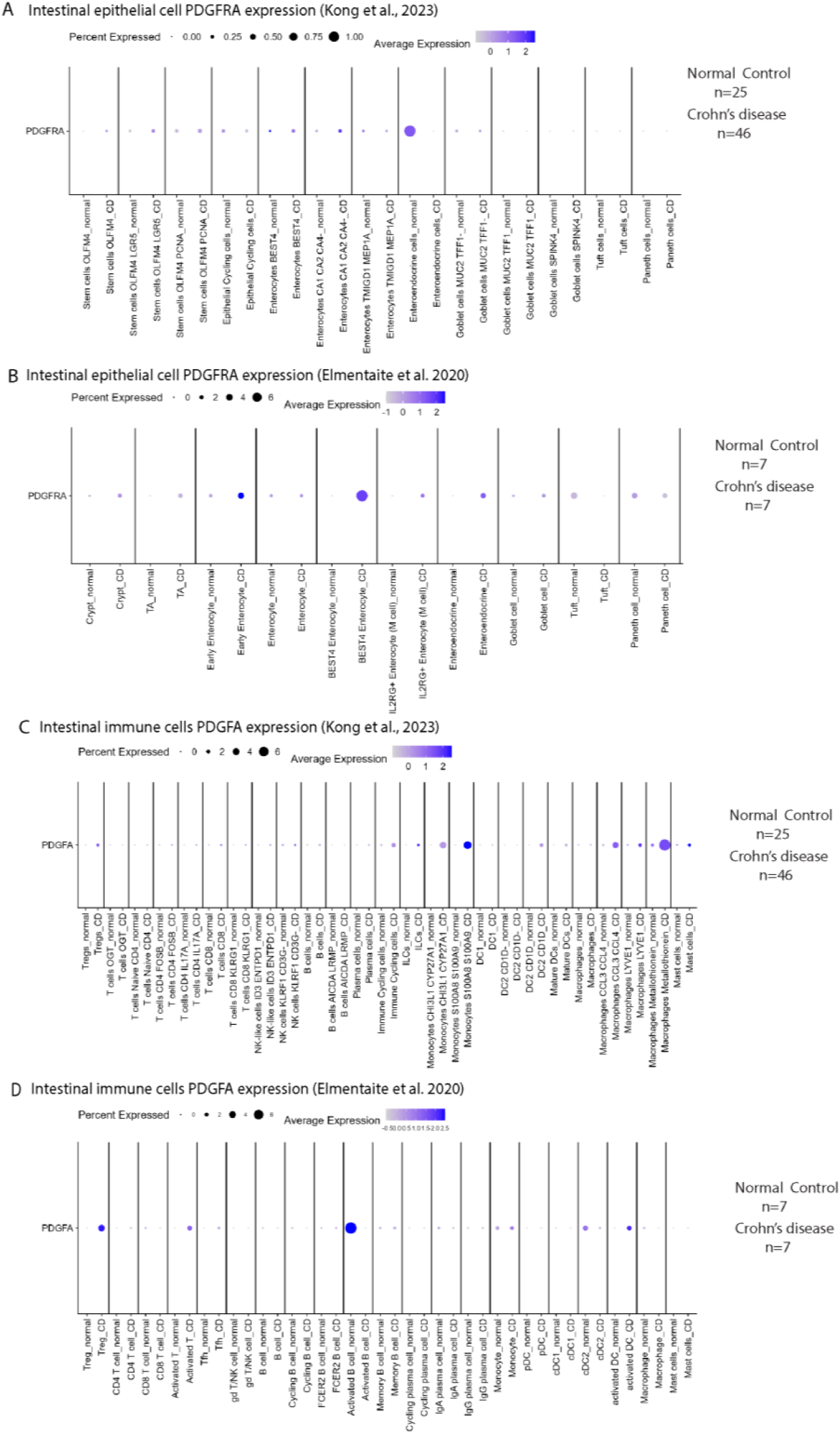
PDGFRA and PDGF-AA expression differ across epithelial and immune cell subsets in normal and Crohn’s disease intestinal tissue. **(A, B)** Dot plots showing PDGFRA gene expression across distinct epithelial cell types in normal donors and CD patients, re-analyzed from study 1 (A) and study 2 (B). **(C, D)** Dot plots showing PDGFA gene expression across distinct immune cell populations in normal donors and CD patients, re-analyzed from study 1 (C) and study 2 (D).

## Supplementary Methods

### Generation of CD4^LVFOXP3^

CD4^LVFOXP3^ were generated as described previously (*12*) fig. S1A. Briefly, peripheral blood was collected in EDTA-coated tubes, and PBMCs were isolated via density gradient centrifugation using Ficoll-Paque. CD4^+^ T cells were subsequently enriched using the CD4^+^ T Cell Isolation Kit (Miltenyi Biotec). The isolated CD4^+^ T cells were activated overnight using TransAct in X-VIVO 15 Medium supplemented with 5% human serum, 100 U/mL IL-2, and 10 ng/mL IL-7. On the following day, activated CD4^+^ T cells were transduced with either LVFOXP3_NGFR or a control NGFR lentivirus at a multiplicity of infection (MOI) of 20, or left untransduced (UT) and cells were maintained in culture with media changes every 2-3 days. On day 5 or 6, transduction efficiency was evaluated by flow cytometry for NGFR expression. On Day 8, NGFR^+^ cells were isolated using the MACSelect NGFR System (Miltenyi Biotec). The enrichment of NGFR^+^ CD4^+^ T cells co-expressing FOXP3 was confirmed on Day 10 through immunostaining for FOXP3 and NGFR, followed by flow cytometry. On Day 11, the transduced cells were re-stimulated and cultured until Day 18 in X-VIVO 15 Medium (Lonza Bioscience) containing 5% human serum, 100 U/mL IL-2, and 1 ng/mL IL-15. On day 18, cells were assessed for NGFR and FOXP3 expression and cryopreserved until use. Activated CD4^+^ T(CD4act) cells or CD4^+^ T cells transduced with the lentivirus containing the NGFR gene only were used as controls for CD4^LVFOXP3^.

### Isolation of tT_reg_ cells

PBMCs were isolated from a leukocyte-rich solution (LRS) chamber and CD4^+^CD25^+^ T-cells were enriched using a CD4^+^CD25^+^ Regulatory T Cell Isolation Kit. The isolated CD4^+^CD25^+^ cells were then stained with antibodies against CD3 (1:50), CD4 (1:50), CD25 (1:50), and CD127 (1:50) and CD14 (1:50) along with a viability dye to exclude dead cells. After staining, the cells were sorted

on a BD FACSAria™ III flow cytometer to obtain a pure population of tT_reg_ (CD4^+^ CD25^high^CD127^low^). The sorted tT_reg_ were washed with X-VIVO 15 Medium containing 5% human serum twice and subsequently activated with 1:1 Dynabeads™ Human T-Activator CD3/CD28 in X-VIVO 15 Medium supplemented with 300 U/mL IL-2 for 10 days. CD4^+^ CD25^-^ T cells were also activated using 1:10 Dynabeads™ Human T-Activator CD3/CD28. Activated CD4^+^ CD25^-^(CD4act) T cells were used as a control for tT_reg_ cells. After 10 days, the Dynabeads were removed, and the activated tT_reg_ and CD4act were cryopreserved until further use in co-culture experiments. For tT_reg_-specific demethylated region (TSDR) analysis, cell pellets were thawed, and genomic DNA was isolated using Dynabeads™ SILANE Genomic DNA Kit. Bisulfite conversion of the extracted DNA was performed to enable the analysis of methylation status and was analyzed using qPCR as described previously (*50*).

### Suppression assay

For the suppression assay, allogenic responder CD4^+^ T cells (50,000 cells) were purified from HD PBMCs using the CD4^+^ T Cell Isolation Kit (Miltenyi Biotec). CD4^+^ responder cells were labeled with 1μM CFSE, and suppressor CD4^LVFOXP3^ or CD4^NGFR^ (25,000–50,000 cells) were labeled with 5μM CellTrace Violet (Fisher Scientific) as per the manufacturer’s instructions in the dark at room temperature. Responders and suppressors at different concentrations (1:0.5–1:1) were activated with Dynabead Human T cell Activator CD3/CD28 at 1:25 bead: cell ratio and co-cultured for 96 hours. 96 hours after stimulation, the cells were immunostained with an antibody to CD4, CD3, and Live/Dead dye, and dilution of CFSE in responder CFSE^+^ live CD3^+^ CD4^+^ T cells was assessed by Flow cytometry as described above. Suppression Index was calculated as follows : Suppression Index = % proliferation (responder + responder) - % proliferation (responder + responder)−% proliferation (responder + CD4LVFOXP3)×100

### PDGF-AA signaling

Single-cell suspensions of enteroid cells were stimulated with recombinant PDGF-AA (50 ng/mL) for 15 minutes and fixation was performed by adding Fix/Perm buffer (1:1, v/v; Foxp3/Transcription Factor Staining Buffer Set). Cells were then permeabilized overnight in cold True-Phos™ Perm Buffer at -20 °C. The next day, cells were washed and stained with antibodies to pAKT (S473), pERK1/2 (Thr202/Tyr204) and LGR5 for 30 minutes in the dark at room temperature, then analyzed by flow cytometry on a BD FACSymphony™ A5. Data were analyzed using FlowJo software.

### Flow cytometry

For tT_reg_-associated marker phenotyping, CD4^LVFOXP3^, CD4^NGFR^ and tT_reg_ cells were cultured for 48 hours in the absence of cytokines. Cells were collected, washed in PBS and stained with Live/Dead Fixable viability dye together with surface antibodies in FACS buffer (PBS, 2% FBS, 2 mM EDTA) for 30 minutes at 4°C in the dark in the presence of Fc receptor–blocking reagent (Miltenyi Biotec). Surface panel included CD3 (1:50), CD4 (1:50), NGFR (1:50), IL1R (1:20), IL6R (1:20), TIGIT (1:20), GITR (1:20), PD1 (1:20), CD127 (1:20) and CD25 (1:20). Cells were washed, then fixed and permeabilized using the Foxp3/Transcription Factor Staining Buffer Set (Invitrogen). Intracellular staining for HELIOS (1:20), FOXP3 (1:20) and CTLA4 (1:20) was performed for 30 minutes at room temperature in the dark.

For enteroid–CD4 T cell co-culture analysis, samples were harvested and processed into single-cell suspensions as described above, then stained with Live/Dead Fixable viability dye and surface antibodies to CD3 (1:50) and CD4 (1:50) or PDGFRα (1:50), followed by intracellular staining for LGR5 (1:20) as described above. Cells were washed, resuspended in FACS buffer, and acquired on a FACSymphony Cell Analyzer; data were analyzed using FlowJo software. Details of the antibodies are provided in the Supplementary Table 2.

### Western blot analysis

0.25 x10^6^ cells were lysed in SDS lysis buffer containing (10% glycerol, 2% SDS, 200 mM DTT, 0.002% bromophenol blue, and 50 mM Tris-Cl pH 6.8) and boiled at 98 °C for 10 minutes. Cells lysed were separated on a 11% gel and transferred to a PVDF membrane and incubated with primary antibodies at the following concentrations: pAKT (1:1000) in 1% BSA in 1X Tris-buffered saline–Tween 20 (TBST) or PDGFRα (1:1000), AKT (1:1000), and HRP-tagged anti-β-actin antibody (1:2500 dilution) in 5% skimmed milk overnight at 4°C on rocker. The next day, the membranes were washed thrice with TBST and incubated with horseradish peroxidase–conjugated secondary antibody (1:1000 dilution) for 1 hour at ambient temperature. Membranes were developed using Super Signal West Dura substrate, and images were acquired using Biorad’s ChemiDoc XRS+ Imaging System. Quantification of western blot was performed using FIJI software.

---

**Supplementary Table 1:**
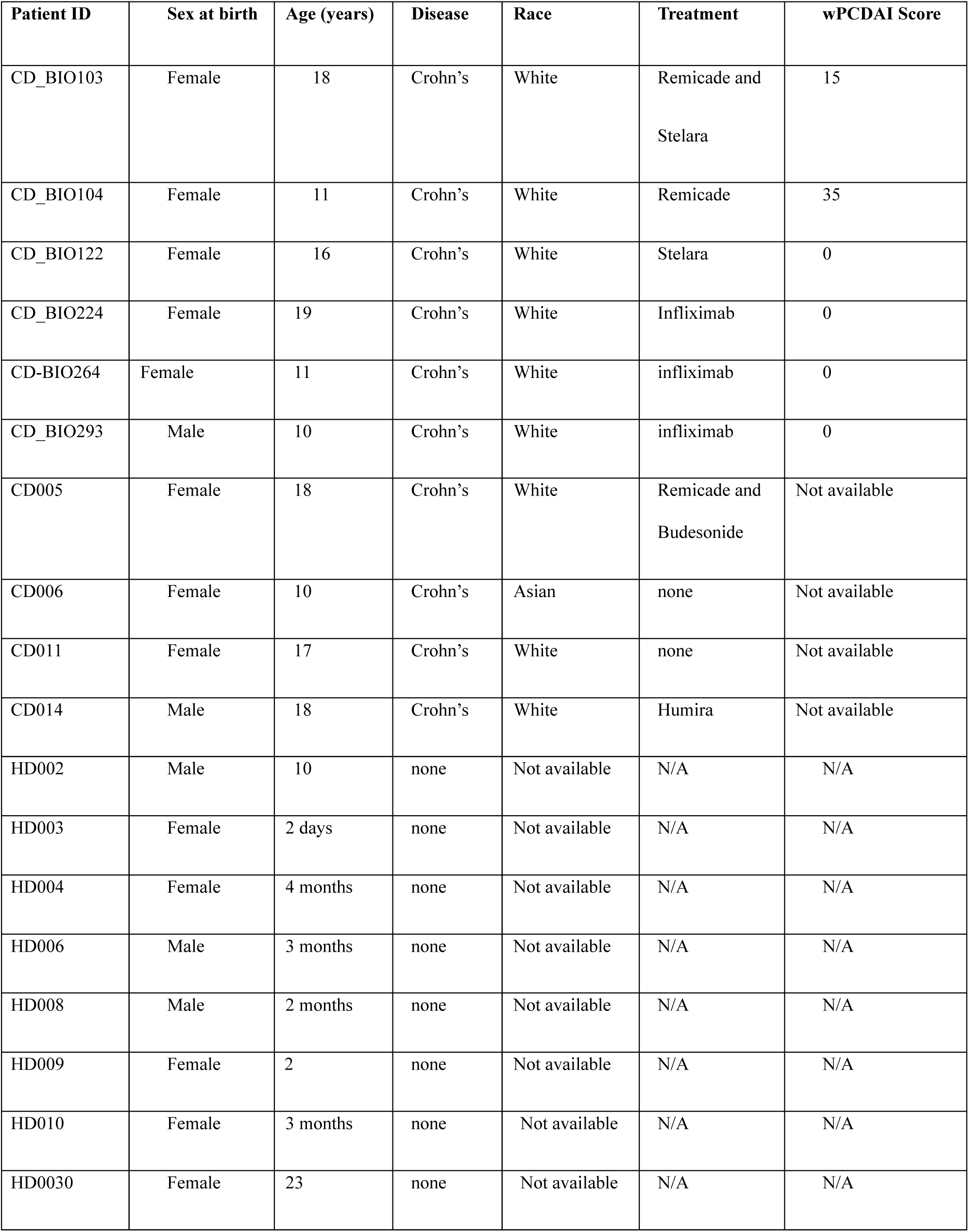
Patient Demographics.

**Supplementary Table 2:** List of antibodies.

| **Antibody target** | **Company** | **Clone** | **Catalogue** | **RRID** |
| --- | --- | --- | --- | --- |
| NGFR (CD271) | BioLegend | ME20.4 | 345108 | AB_10645515 |
| FOXP3 | BioLegend | 259D | 320212 | AB_430887 |
| CD3 | BioLegend | OKT3 | 317336 | AB_2561628 |
| CD4 | BioLegend | RPAT4 | 300518 | AB_314086 |
| IL6R | BioLegend | UV4 | 352812 | AB_2562719 |
| ICOS | BioLegend | C398.4A | 313525 | AB_2562642 |
| TIGIT | BioLegend | A15153G | 372735 | AB_2892455 |
| CTLA4 | BioLegend | BNI3 | 369631 | AB_2892450 |
| PD1 | BioLegend | NAT105 | 367430 | AB_2734414 |
| CD127 | BioLegend | A019D5 | 351310 | AB_10960140 |
| CD14 | BioLegend | M5E2 | 301828 | AB_2275670 |
| HELIOS | BioLegend | 22F6 | 137232 | AB_2565797 |
| GITR | BioLegend | 621 | 311610 | AB_2204247 |
| pERK1/2(Thr202/Tyr204) | BioLegend | 6B8B69 | 369519 | AB_2721567 |
| pAKT (S473) | BioLegend | A21001C | 606555 | AB_3068117 |
| PDGFRα | BioLegend | 16A1 | 323511 | AB_2783190 |
| CD25 | BD Biosciences | 2A3 | 562660 | AB_2744343 |
| LGR5 | R&D Systems | 707042 | FAB8078P | AB_3652859 |
| IL1R | R&D Systems | Polyclonal | FAB269P | AB_2124912 |
| PDGFRα | Abcam | EPR22059-270 | ab203491 | AB_2892065 |
| β-catenin | Abcam | E247 | ab32572 | AB_725966 |
| β-actin | CST | 8H10D10 | 12262S | AB_2566811 |
| pAKT (S473) | CST | 587F11 | 4051S | AB_331158 |
| AKT | CST | Polyclonal | 9272S | AB_329827 |
| Anti-PDGF-AA | R&D Systems | 114506 | MAB221-100 | AB_2252175 |
| Mouse IgG1 Isotype control | Bio X Cell | MOPC-21 | BE0083 | AB_1107784 |

## Table of Reagents

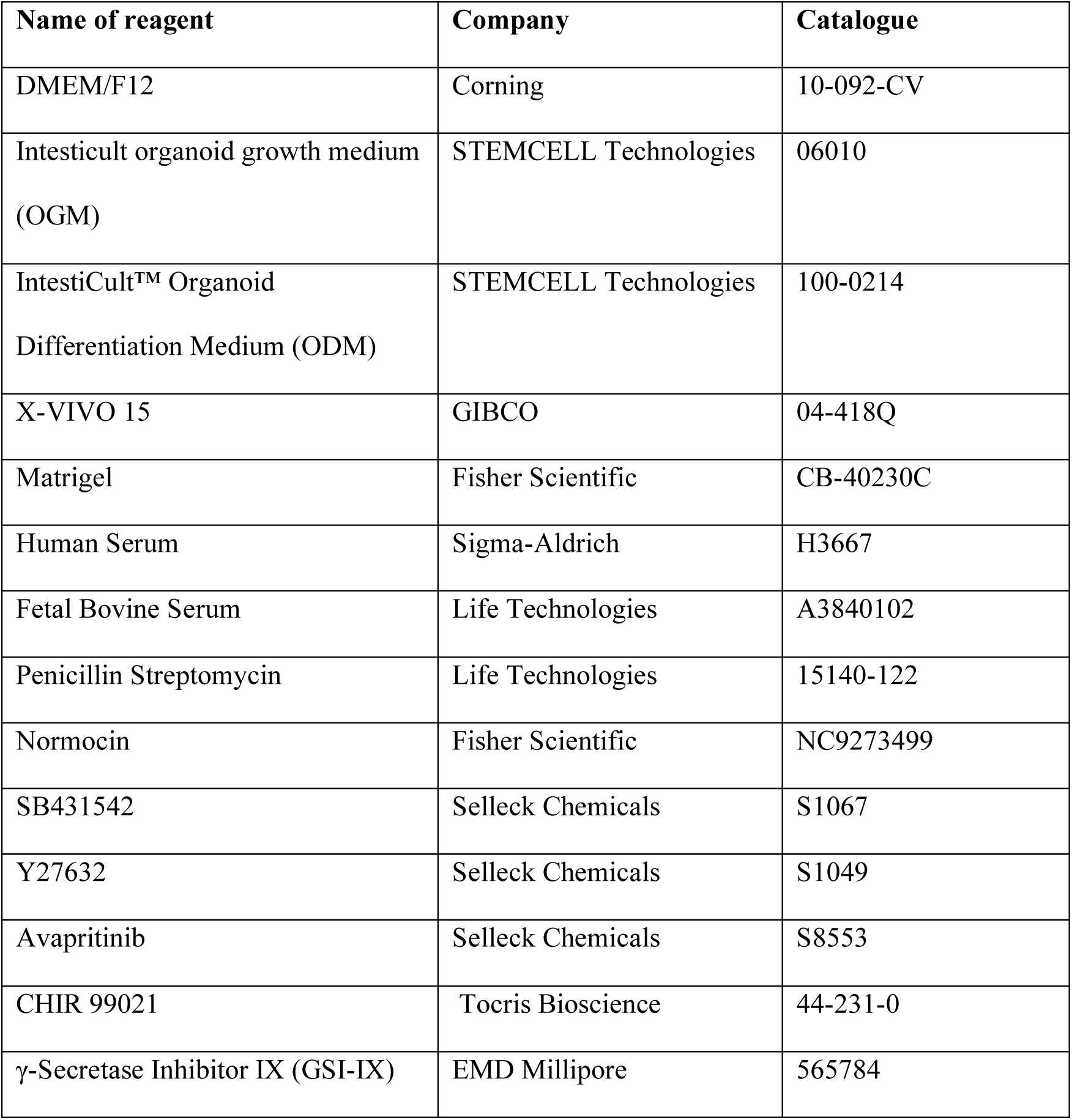
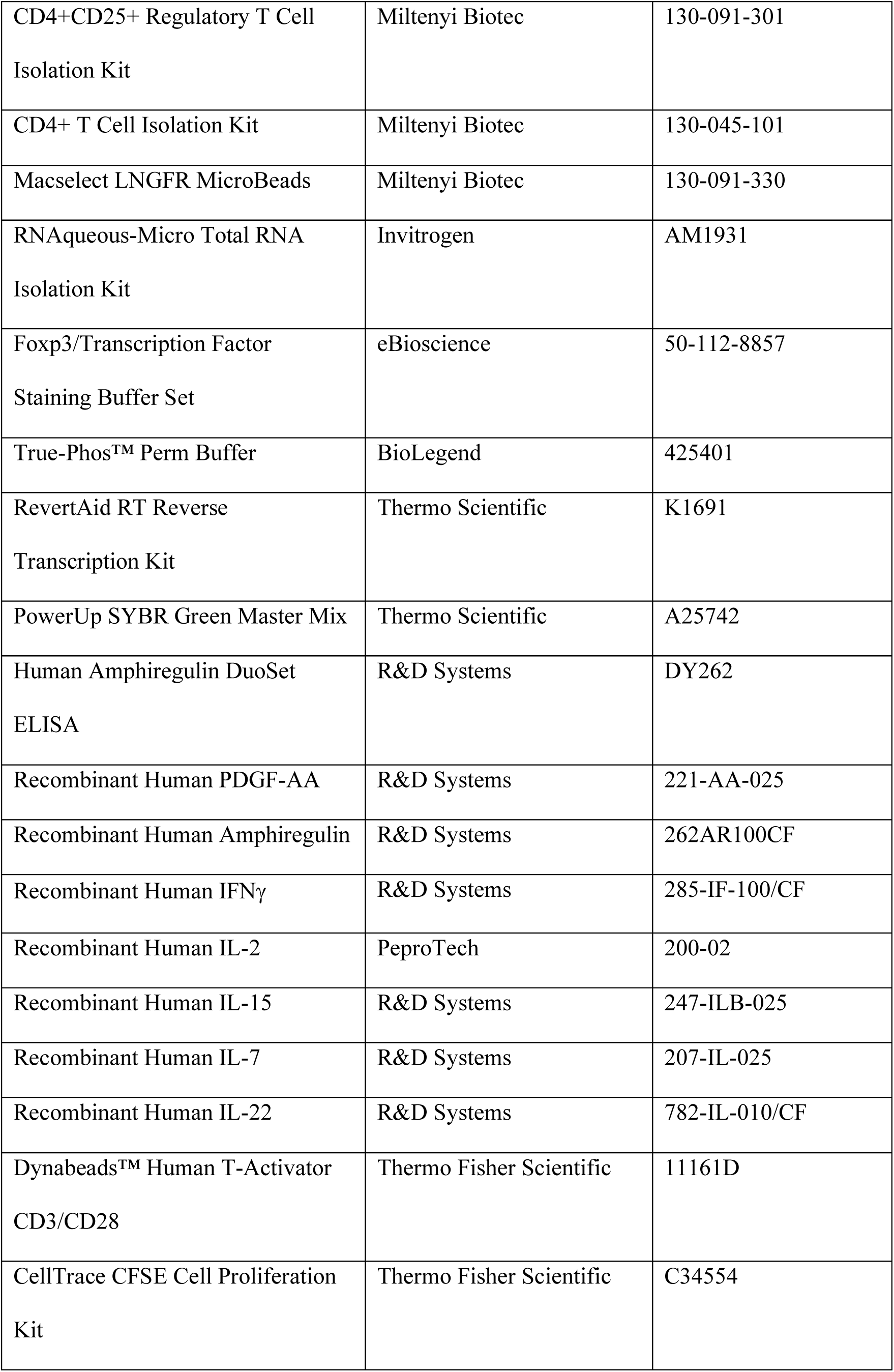
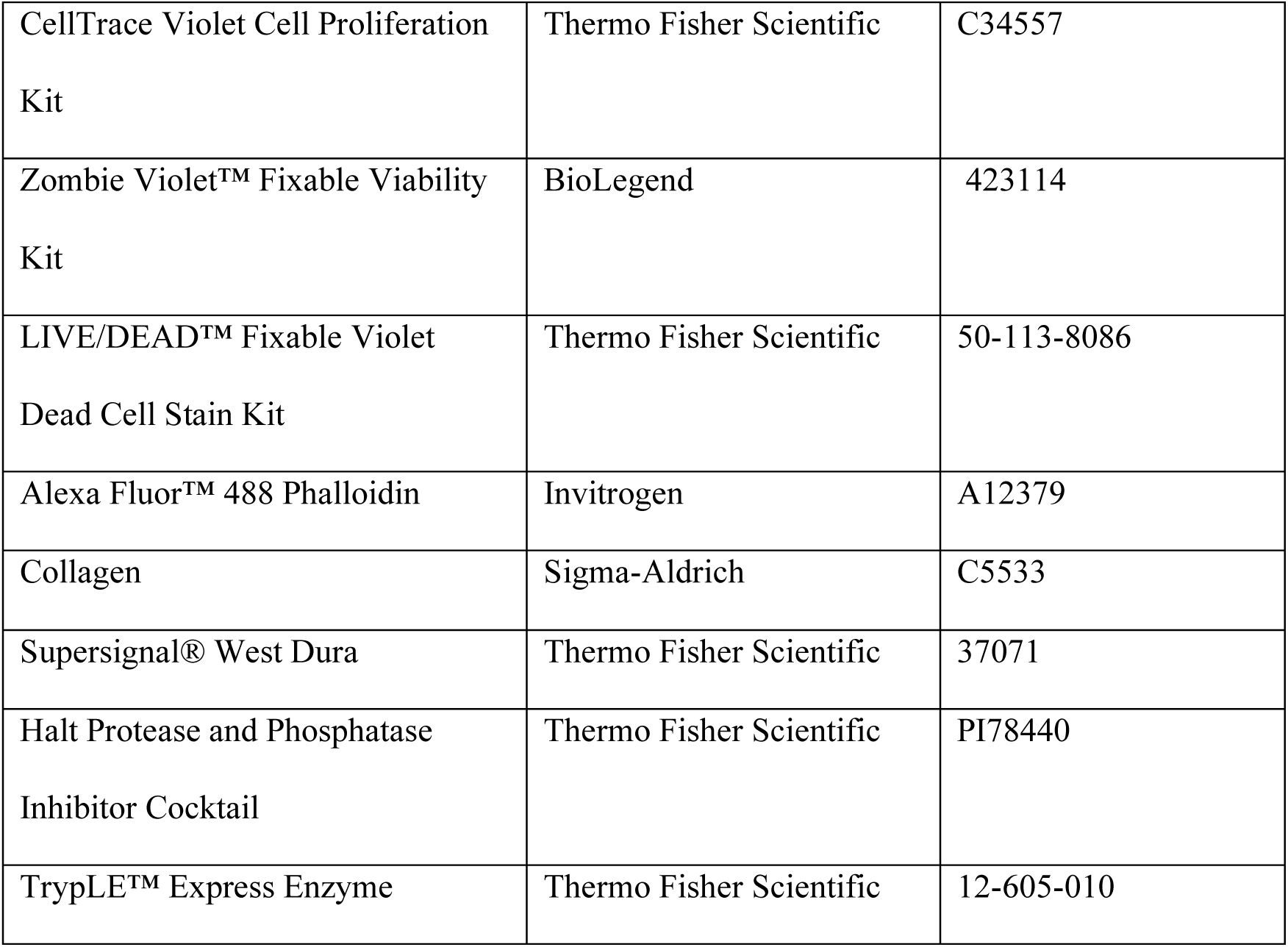

